# Juggling the Limits of Lucidity: Searching for Cognitive Constraints in Dream Motor Practice

**DOI:** 10.1101/2025.02.12.637898

**Authors:** Emma Peters, Kathrin Fischer, Daniel Erlacher

## Abstract

Lucid dreaming (LD), during which the dreamer becomes aware of the dream state, offers a unique opportunity for a variety of applications, including motor practice, personal well-being, and nightmare therapy. However, these applications largely depend on a dreamer’s ability to control their dreams. While LD research has traditionally focused on induction techniques to increase dream frequency, the equally crucial skill of dream control remains underexplored. This study provides an initial investigation into the mechanisms of dream control and its potential influencing factors. We specifically examined whether a complex motor skill—juggling—could be performed within a lucid dream, creating a particularly challenging lucid dream task, which calls for a high level of dream control. Eight healthy participants (aged 24–50) underwent overnight polysomnography (PSG) at the University of Bern’s Institute for Sports Science, provided detailed dream reports, and completed questionnaires assessing dream control, self-efficacy, personality traits, mindfulness, motivation, and intention setting. Of these, four participants experienced lucid dreams, and of these, two demonstrated high dream control with successful LD juggling attempts. Trait differences between non-lucid and lucid dreamers in the lab were examined, with a focus on low-to-no dream control versus high dream control among the lucid dreamers. The two lucid dream juggling attempts are described in detail, providing insight into the challenges of executing complex tasks within a lucid dream. While this study lacks in sample size, it highlights the potential roles of many psychological traits, such as belief, motivation, and self-efficacy, in shaping dream control abilities. This study helps to lay the groundwork for future research aimed at investigating lucid dream control and therefore optimizing LD applications in therapy, sports training, and cognitive science.

## 1. Introduction

### 1.1 Wakeful mental rehearsal of motor skills as the foundation for lucid dream motor practice

People often assume that improving physical skills requires repeated physical practice. However, this is not necessarily the case. Mental practice is a well-established method for improving motor skills without physical movement. It involves the mental rehearsal of a task, engaging neural circuits similar to those activated during actual execution (Feltz et al., 1988; Jeannerod, 2001). According to neural simulation theory, motor imagery strengthens pathways involved in movement control, enhancing skill acquisition (Lotze & Halsband, 2006; Munzert et al., 2009). Research shows that mental practice benefits both sports and rehabilitation. It has been found to enhance fine motor skills (Guillot et al., 2012), improve gymnastics performance (Cumming & Ste-Marie, 2001), and aid in stroke recovery (Zimmermann-Schlatter et al., 2008). Mental imagery can also help reduce pain and improve function in conditions like chronic back pain (Braun et al., 2019). The effectiveness of mental practice depends on task complexity, imagery vividness, and training frequency (Lebon et al., 2012). There is also a distinction between internal imagery—where one imagines performing an action from a first-person perspective—and external imagery, where one observes themselves from an outside view. Internal imagery tends to be more effective for precise, goal-directed tasks, whereas external imagery is better suited for activities requiring adaptability (Lebon et al., 2012). Although mental practice alone can improve performance, combining it with physical practice often leads to greater benefits than either approach alone (Feltz et al., 1988; McBride & Rothstein, 1979). This combination makes it a valuable tool in sports training, motor learning, and rehabilitation.

### 1.2 Lucid dreaming as an immersive, virtual reality-like practice environment

Building on the concept of mental practice, lucid dreaming offers a unique extension for motor learning by allowing movements to be rehearsed within dreams. Lucid dreaming occurs when individuals become aware that they are dreaming and, in some cases, gain control over the dream environment (Baird et al., 2019). This hybrid state of wake-like cognition within REM sleep (Hobson, 2009) provides an opportunity to engage in mental and motor imagery with heightened immersion. Similar to Jeannerod’s neural simulation theory (Jeannerod, 2001), lucid dreaming can be classified as an “s-state,” in which imagined and executed movements activate overlapping sensorimotor regions. An fMRI and near-infrared spectroscopy study showed that motor tasks performed in lucid dreams engage the same neural structures as waking actions (Dresler et al., 2012). This supports the idea that motor rehearsal in dreams may strengthen neural pathways, just as mental practice does in wakefulness. Physiological markers further confirm the connection between dreamed and real movements. Using electrooculography (EOG), voluntary eye movements during dreams can be recorded, enabling objective verification of the lucid dream (LaBerge, 1990). Studies using electromyography (EMG) have also detected muscle twitches corresponding to dreamt movements, despite REM-related muscle atonia (Fenwick et al., 1984; LaBerge et al., 1981). These findings suggest that dreamed motor actions generate real neuromuscular activity, strengthening the potential of lucid dreaming as a motor training tool. Further research highlights that movement planning in lucid dreams resembles waking motor control. For example, sensorimotor area (SMA) activation during dreamt hand movements was found (Erlacher et al., 2003), as well as increased SMA activation in lucid dreamers performing imagined movements, comparable to actual execution (Dresler et al., 2011). Since the SMA plays a key role in movement preparation, these results suggest that lucid dream practice could enhance real-world motor skills by fine tuning the movement preparation. While promising, lucid dream motor practice faces challenges. Dreamers often struggle with maintaining stability, controlling dream environments, or sustaining lucidity (Schredl & Erlacher, 2011). However, if lucid dream control can be improved, motor learning within dreams could provide a powerful adjunct to mental and physical practice, benefiting areas such as rehabilitation, skill acquisition, and performance enhancement.

### 1.3 Lucid Dream Motor Practice can improve performance during wakefulness

Research on motor performance enhancement through lucid dreaming suggests that practicing movements in dreams can be a valuable tool for improving waking task performance and overall wellbeing. In addition to motor skill development, lucid dreaming has been associated with enhanced problem-solving abilities, creativity, and emotional regulation (Schädlich & Erlacher, 2012). Understanding the neural mechanisms behind lucid dreaming could have significant implications for cognitive and clinical settings, including the development of novel treatments for nightmare disorders and cognitive performance enhancement (Mota-Rolim & Araujo, 2013). A meta-analysis by Bonamino et al. (Bonamino et al., 2023) provides strong evidence that lucid dream practice can positively impact waking task performance. The findings suggest that individuals who engage in lucid dream practice may experience improvements in cognitive function and problem-solving skills, ultimately leading to enhanced performance in various waking-life activities. The study also highlights potential benefits for individuals with sleep disorders, anxiety, depression, and those seeking to improve athletic or academic performance (Bonamino et al., 2023). Similarly, a review by Peters et al. (Peters et al., 2023) supports the idea that lucid dreaming can be an effective tool for enhancing motor skills. The study suggests that practicing movements in lucid dreams may lead to measurable improvements in activities such as playing an instrument or engaging in sports. This is particularly valuable for individuals who are unable to practice physical activities in waking life due to injury or other constraints. By allowing for continued motor skill rehearsal during sleep, lucid dreaming can create better performance during wakefulness. The review also emphasizes that lucid dreaming has the potential to enhance not only motor abilities but also cognitive functions and emotional resilience. Moreover, lucid dreaming provides a unique advantage over traditional mental practice by offering an immersive, virtual reality-like experience.

### 1.4 Lucid dream motor practice efficacy depends on amount of dream control

Motor practice in lucid dreams is a form of mental rehearsal, allowing dreamers to refine motor skills while physically asleep. Studies support its effectiveness in enhancing motor performance (Stumbrys et al., 2016). In a study on finger-tapping, Stumbrys et al. (2016) compared lucid dream practice with physical and mental practice in wakefulness and a control group without any practice. Results demonstrated significant improvement in all practice groups, supporting the notion that lucid dream motor practice is comparable to both physical and mental rehearsal. Notably, engaging in motor practice during sleep did not negatively affect subjective sleep quality. Similarly, Erlacher and Schredl (Erlacher & Schredl, 2010) examined motor practice in lucid dreams using a coin-tossing task. Participants attempted to toss a 10-cent coin into a cup, aiming for the highest possible success rate across 20 tosses. While physical practice showed the greatest improvement, lucid dream practice also led to enhanced performance upon waking. However, it remains unclear whether this improvement resulted solely from dream practice or if other contributing factors played a role. Further investigation explored how distractions influence motor practice within lucid dreams (Schädlich et al., 2017). In a dart-throwing task, researchers observed that both the level of dream control and the presence of distractions affected performance outcomes. One participant reported interference from a dream character: “The doll kept throwing darts at me.” Another noted instability in the dream environment: “I noticed it was getting somewhat unstable… I performed another eye signal… I managed three or four more throws and then I woke up” (Schädlich et al., 2017). These findings suggest that much like in wakefulness, distractions in the dream world can hinder us during task rehearsal. Research on waking-life motor learning supports this complexity. Beilock and Carr (Beilock & Carr, 2001) found that distractions during juggling practice led to slower learning and reduced accuracy. Conversely, Maslovat et al. (Maslovat et al., 2010) showed that distractions requiring cognitive processing similar to the primary motor task could actually improve motor learning. Thus, the impact of distractions on both wakeful and lucid dream practice appears to depend on their characteristics. While lucid dreaming offers significant potential for motor training, there are limitations to its applicability. Not all individuals can reliably achieve lucidity, and those who do may struggle to maintain the state long enough to engage in sustained practice (Schredl & Erlacher, 2011). There is a wide variability in lucidity between and within dreams and dreamers (Mallett et al., 2021) and this includes the ability to control aspects of the dream. This raises the question of whether certain personality traits or specific cognitive abilities influence dream control and whether these abilities can be cultivated through training. The present study aims to deepen our understanding of the challenges and constraints associated with motor practice in lucid dreams, particularly in relation to the role of distractions. In a lucid dream, individuals have the potential to exert full control over their surroundings, bodily sensations, and dream characters, creating a highly immersive training environment (Schredl & Erlacher, 2008). Given the parallels between the dreaming and waking body, lucid dream practice could serve as a valuable tool for refining real-world motor skills. However, achieving a stable and controlled dream state remains essential for maximizing its benefits in skill improvement, but also clinical applications such as lucid dream nightmare therapy (Ouchene et al., 2023).

### 1.5 The level of dream control might depend on various cognitive factors

While lucid dreaming has potential for motor practice, a major challenge lies in the lack of control over the dream environment and the unpredictability of dream scenarios. The ability to manipulate dream elements and perform complex tasks varies widely among individuals due to differences in cognitive abilities, dream recall, and dream stability (Schredl & Erlacher, 2011; Tholey, 1983). Some lucid dreamers can engage in advanced activities like solving mathematical problems, playing instruments, or juggling, whereas others struggle to exert even basic control over their dreams. A study by Schredl et al. (Schredl et al., 2018) highlights this variability. Among a group of lucid dreamers, 42% reported at least one lucid dream per month, but only 11% consistently managed to alter three-quarters or more of their lucid dreams. Furthermore, about half of all attempts to change specific aspects of lucid dreams ended in failure (Stumbrys et al., 2014). This distinction between lucid insight (awareness of dreaming) and control or manipulation (modifying the dream environment) suggests that awareness alone does not guarantee the ability to alter the dream world. Lemyre (Lemyre et al., 2020) explored dream control by creating five categories of Lucid Dream Control Strategies (LDCS); verbal strategies, strategies based on the use of the dream’s objects or environment, strategies based on the use of the oneiric body, strategies based on the management of emotions, and other strategies. This categorization helped to identify different LDCS that were used individually, or in some type of combination.

Dream control challenges are further evident in lucid nightmares, where dreamers remain aware but experience reduced agency and emotional distress (McNamara et al., 2015). Common features include feelings of helplessness, aggressive dream characters, and difficulties in waking up. Research indicates that lucid nightmares occur more frequently in women, individuals with a history of nightmares, and spontaneous lucid dreamers rather than those who induce lucidity intentionally (Stumbrys, 2018). Given these findings, controlling lucid nightmares may be even more difficult than controlling regular lucid dreams.

The question remains whether dream control is a trainable skill that can be systematically developed or if it is inherently constrained by individual differences in cognitive and personality traits. One potential factor influencing dream control is self-efficacy, which refers to an individual’s belief in their ability to achieve specific goals (Bandura, 1997). Just as self-efficacy varies across domains—such as sports, academics, or social interactions—its influence on dream control remains uncertain. While some studies suggest that higher self-confidence correlates with frequent lucid dreaming (Doll et al., 2009), no direct evidence yet confirms a causal relationship between self-efficacy and dream control. Research has linked frequent lucid dreaming to various mental health and personality traits. Doll et al. (2009) found that frequent lucid dreamers exhibited greater assertiveness, emotional endurance, and life satisfaction, as well as higher dream recall frequency. Similarly, studies by Schredl et al. (Schredl et al., 1996) and Cernovsky (Cernovsky, 1984) suggest that a positive attitude toward dreams correlates with higher dream recall. Moreover, openness to experience, creativity, and alexithymia have been associated with both increased dream recall and heightened dream insight (Brand et al., 2011; Ruby, 2011; Schredl & Erlacher, 2007). However, these findings do not necessarily indicate better dream control. While frequent lucid dreamers recall dreams more vividly, there is no conclusive evidence that they have greater control over dream content than rare lucid dreamers. Yet, many lucid dreamers report difficulty carrying out intended tasks due to insufficient clarity or fading lucidity (Stumbrys et al., 2014). Lucidity can even diminish mid-task, or dreamers may complete a task without fully realizing they are dreaming (Worsley, 1984). Moss (Moss, 1986) introduced the Dream Lucidity Continuum, which suggests that lucidity exists on a spectrum, ranging from minimal dream awareness to full control. This lucid spectrum was backed by Mallett (Mallett et al., 2021), who further investigated the variability of lucidity between and within individual. However, the factors that facilitate higher control and sustained dream clarity remain poorly understood. Despite the evidence linking frequent lucid dreams to higher dream recall and metacognitive awareness, there is no established link between lucid dream frequency and dream control. This raises an important question: Do the same factors that enhance lucid dream frequency also improve dream control? Additionally, if dream control is a trainable skill, could it be developed through deliberate practice, much like lucid dreaming itself? One study used online dream reports and questionnaires to investigate the relationship between dream control and several of the factors mentioned above. Lucid dreaming frequency and previous knowledge seem to be the main predictors of higher lucid dream skills (Sammer et al., 2024). While other factors, including self-efficacy, beliefs about LD, and meta-awareness were investigated, they did not predict more dream control. More research is needed to determine which variables optimize dream control. A better understanding of these factors could enhance the feasibility of motor practice in lucid dreams and other applications of lucid dreaming, such as lucid dream therapy.

This work is a further exploration on influencing and controlling dream content and performing complex motor tasks in a state of lucid dreaming in the sleep lab. Therefore, motor skills and control over different dream aspects in lucid dreams are examined and compared. Moreover, different variables are questioned with an additional online survey such as motivation and persistence, self-efficacy, personal believes and attitude, stress level and mindfulness, which may contribute to a higher level of dream control. The choice of juggling as the motor task is motivated by its high complexity. It demands many different skills and factors like executing controlled movements and coordination, additional material or equipment, and earthly environmental factors such as gravity. The aim is to provoke as much complexity and distractions as possible in order to visualize and explore them.

## 2. Methods

### 2.1 Study design

This study invited experienced lucid dreamers to spend a night in the sleep laboratory at the University of Bern to practice a motor task within a lucid dream. A qualitative analysis of different dream reports was conducted based on the experiment, alongside surveys assessing overall sleep habits, personality traits, and study-specific questions. Participants who demonstrated success or showed promising signs but were unable to complete the experiment in a single session were invited for multiple sessions to collect additional data on the motor task in a lucid dream. The primary objective was to investigate the feasibility of a complex motor task – juggling-during lucid dreaming and explore potential cognitive parameters which might influence its success.

### 2.2 Participant recruitment and selection

A total of eight healthy individuals (N female = 2 age: 24-50) participated in this study. All participants provided informed consent and were fully briefed on the study protocol, including their right to withdraw at any time. Ethical approval was obtained, with no further concerns raised. The participants spent in total eleven nights in the sleep laboratory, with participants spending between one and three nights in the sleep lab. Most participants (N=6) underwent a single night of observation and measurement, while two were invited back for additional sessions due to challenges encountered or exceptionally promising initial performance.

### 2.3 Experimental setting and equipment

*Sleep laboratory -* The study took place in the sleep lab at the Institute for Sports Science, University of Bern. Each participant spent the night in a bedroom equipped with a bed, a surveillance camera for observation, a microphone for communication with the experimenter in the control room, and the necessary equipment for polysomnography (PSG).

*Polysomnography -* Participants were monitored using a 10-channel PSG setup, including EEG (F3, F4, C3, C4, O1, O2), horizontal EOG, EMG (chin), a reference electrode (Fz), and a ground electrode (A2, right earlobe). Electrodes were connected to a gUSBamp amplifier, following the standardized 10-20 EEG system (Homan et al., 1987). Before sleep, the pre-agreed eye signal [left-right-left-right]) was rehearsed to ensure signal accuracy and facilitate later recognition.

### 2.4 Questionnaires

Participants were first asked for demographic data including names, age, gender, and occupation. To assess more information about the participants general experiences and skills in lucid dreaming as well as personality traits, an additional extensive questionnaire was created specifically designed for this study. This questionnaire includes sections combined from pre-existing questionnaires and additional study-specific questions were incorporated. The questionnaire was formulated using the LimeSurvey online survey tool. To ensure uniform comparability, the survey was conducted in English.

The questionnaire included the following components:

1. Juggling skills
2. Dream recall
3. Lucid dream experience
4. Induction techniques
5. Reality testing
6. Dream goals
7. Dream control
8. Motivation and persistence
9. Self-efficacy
10. Personal beliefs and attitudes
11. Stress levels
12. Mindfulness

In the following sections a more detailed description of the various components of the questionnaire is provided.

#### 2.4.1 Juggling skills

Participants were asked about their skills in juggling, specifically assessing their ability to juggle with three objects. These questions aimed to see any potential differences in juggling skills between waking life and lucid dreaming. Participants were further asked to indicate their level of confidence in juggling during wakefulness on a 5-point scale with 1 meaning “not successful” and 5 meaning “very successful”. These items would later be asked again during the dream report. This might help explore potential variations in motor skill performance and confidence levels between the two states, contributing to understanding the impact of the dreaming state on juggling abilities.

#### 2.4.2 Dream recall

In evaluating the frequency of dream recall among participants, a 7-point Dream Recall Frequency scale developed by Schredl (Schredl, 2004) was employed.

#### 2.4.3 Lucid dreaming experience

To determine the suitability of participants for this study, they were queried about whether they had ever experienced a lucid dream and about the frequency of their lucid dreams, as well as how they normally become lucid.

#### 2.4.4 Induction techniques

Specific induction techniques and their effectiveness were assessed. Participants were presented with different options including the most frequently used techniques in lucid dreaming research. They consist of the following (Reality testing, Noticing the unreal nature of the dream while dreaming (DILD), Intention setting before sleep, Wake-back-to-bed (WBTB), setting an intention to remember to recognize when one is dreaming (MILD), maintaining awareness while transitioning from wakefulness to sleep (WILD), Focusing on sensory stimuli and repeatedly shifting the attention between different sensations (SSILD), Others). For a full review on lucid dream induction strategies see (Tan & Fan, 2022). Participants were also asked how frequently they practice lucid dreaming techniques. It was also assessed if they keep a dream journal to record their dreams and if so, how often they use it in the last few months as well as how effective it has been for improving their lucid dreaming skills.

#### 2.4.5 Reality testing

To assess the use of reality testing, participants were asked whether they regularly perform reality tests during their waking hours in order to check if they are dreaming. They were also inquired about the methods they use for reality testing, with the most popular options such as searching for abnormal things, double-checking the time, trying to push a finger through the palm of the opposite hand, checking their reflection, looking at their hands, reading, and attempting to switch a light on/off. For each method, participants were requested to rate its effectiveness on a scale from 1 to 5, where 1 means “not effective at all,” and 5 means “extremely effective.”

#### 2.4.6 Dream goals

Participants were asked if they set specific goals or intentions for their dreams, and additionally asked to rate their success in achieving these dream goals on a scale ranging from 1 to 5. On this scale, a rating of 1 indicated “not successful at all,” 5 meaning “extremely successful,” and there was an option for participants who did not set any goals.

#### 2.4.7 Dream control

Participants were presented with a set of questions assessing their experiences and capabilities related to dream control. Typical behaviors after achieving lucidity were assessed, with options including changing the dream scenery, interacting with dream characters, manipulating objects or events, flying or levitating, observing without interference, and an open-ended option for other behaviors. These were rated using a scale from 1 meaning “not successful” to 5 indicating “extremely successful” to measure participants’ perceived ability to control various aspects of their dreams, including elements like dream body, other characters’ bodies, movement, actions, objects, environment, time, and gravity.

Participants were further asked to reflect on challenges or limitations experienced during dream control attempts, addressing aspects such as maintaining lucidity, recalling goals, focus retention, imagining supernatural actions, difficulty distinguishing dreams from reality, and any other challenges they encountered. Specific challenges associated with different parts of dream control, including the dream body, other characters’ bodies, movement, actions, objects, environment, time, and gravity, were examined, with participants providing ratings on a scale from 1 to 5 for each challenge’s impact on dream control.

#### 2.4.8 Motivation and persistence

To assess persistence, a questionnaire from Constantin et al. (Constantin et al., 2011) was used. It consists of 13 questions from which 12 questions were used in this study. Every question had to be answered on a scale from 1 (“very low degree”) to 5 (“very high degree”) with three different subcategories. The questions in the scale of Long-term purposes pursuing reflect if a person likes to keep doing things for a long time, especially bigger projects, showcasing the commitment to set and persevere in achieving future projects. The Current purposes pursuing emphasizes the ability to resist distractions, compensate for challenges and complete tasks once started. Lastly, the Recurrence of unattained purposes reflects the tendency to keep thinking about old projects they didn’t finish or when they become more aware of chances that can help them finish those projects. Furter the participants were asked if they use lucid dreaming for problem solving and what their general reasons or motivations are for lucid dreaming, giving different examples and the option to write others.

#### 2.4.9 Self-efficacy

To evaluate self-efficacy, which refers to their confidence in performing specific actions, questions related to juggling skills were asked. The construction of these questions based on the guidelines provided by Bandura (Bandura, 2005) for constructing self-efficacy scales. They were asked to rate their confidence in successfully executing juggling in a lucid dream on a scale from 0 (“extreme uncertainty”) to 100 (“extreme certainty”). Additionally, participants had to express their confidence in performing juggling in the waking state using the same rating scale.

#### 2.4.10 Personal beliefs and attitudes

Participants’ personal beliefs and attitudes were assessed by asking whether they believed they could become lucid in their dreams. Participants were asked to express their opinions on the potential impact of lucid dreaming on their waking life, asking about their perception of the broader consequences of lucid dream experiences. Additionally, participants were asked if they believed they could engage in juggling during a lucid dream.

#### 2.4.11 Stress levels

The Perceived Stress Scale (PSS), was used to assess stress levels. The scale consists of questions about emotions and thoughts experienced over the past month, rated on a scale from 1 (“never”) to 5 (“very often”). Designed to evaluate perceived stress in terms of unpredictability, uncontrollability, and overload, the PSS also includes direct questions about current stress levels. The results were compared to normative data from a study by Cohen and Janicki-Deverts (Cohen & Janicki-Deverts, 2012).

#### 2.4.12 Mindfulness

To assess the overall level of mindfulness, the Five Facet Mindfulness Questionnaire (FFMQ) from Baer et al. (Baer et al., 2006) was used. This questionnaire uses five distinct parts of mindfulness, Describing, Observing, Acting with Awareness, Nonjudging of inner experience, and Nonreactivity to inner experience. Each part was assessed through two questions, and participants rated their responses on a scale from 1 (“never or very rarely true”) to 5 (“very often or always true.”) Additionally, participants were asked about their engagement in other mindfulness practices, including meditation, yoga, and hypnosis, rating the frequency of these practices on a six-point scale from never to very frequently (more than 5 times per week). The survey also allowed participants to specify any other mindfulness exercises they practice, providing an overview of their mindfulness habits beyond the specified categories.

### 2.5 Procedure and protocol

Before every experiment, the participants received information regarding the goal as well as the walk-through of the study. They were asked to not consume alcohol or caffeine on the day before the experiment. Any open questions or uncertainties were addressed, and a consent form was provided for participants to review and sign.

The experiment started at 21:00 in the evening. As soon as participants were ready for bed, the experimenter applied the PSG. During the application of the PSG, the exact protocol of the night was repeated, and any questions and uncertainties were clarified. Since participants often had a personal strategy or technique for becoming lucid and the primary aim of this study was for the participants to have a lucid dream, the protocol was adapted if necessary. The participants then went to bed with the PSG setup attached and upon identification of N2 sleep stage, a timer was set for approximately four hours, during which participants were able to sleep normally. While REM sleep was detected, the experimenter watched out for the agreed-on eye signal from the participant. In case they became lucid, they were instructed to perform an initial left-right-left-right eye movement (LRLR), followed by an attempt to juggle. Once the participants finished the attempt (as many attempts for as long as they could) they did the same eye movement LRLR once again. After the second signal, they were awakened through the intercom, and a dream report was collected and recorded. In case the participant did not become lucid in the first couple of hours after the initial sleep onset, they were awakened after about four hours and a “wake back to bed” (WBTB) protocol was executed. During the WBTB, the participants were awake in a period of approximately 30-minutes, they engaged in lucid dreaming and sleep-related topics, such as discussing or reading about the topic. They reported their prior dreams and talked about the further steps of their individual method, any uncertainties were clarified, and their goals were for the lucid dream were discussed. Those methods can be highly individualized, as each person has a different technique for becoming lucid (e.g., MILD, WILD, SSILD etc.). After the 30-minute period, the participants could use their individual technique and the experimenter again observed the PSG for the eye signal. If the initial LRLR eye signal was detected, a waiting period began were the experimenter looked for another LRLR signal. If the second LRLR was detected or in case the EEG showed signs of arousal or wakefulness, the participant got called via the intercom system, followed by dream report collection as described above.

### 2.6 Data collection and analysis

Sleep data, including the confirmation of a lucid dream eye-signal were analysed using EEGLAB (Delorme & Makeig, 2004). Questionnaire data was exported from LimeSurvey and handled in Excel. The data in this study is solely reported and described but no statistical tests were performed due to the very small sample size.

Dream reports were collected in German, Swiss German, and English. To ensure consistency and clarity, all reports were standardized following the dream reporting manual by Schredl (Schredl, 2018). Reports originally written in German or Swiss German were then translated into English using the DeepL translation tool. The translated reports were manually reviewed to correct potential inaccuracies and ensure fidelity to the original content.

### 2.7 Ethical consideration

All participants within the study participated in the research on a voluntary basis and were able to cease participation in the experiment at any time. Prior to the study, all participants received information about the study methodology and signed a written consent form. All data collected from the participants were managed with pseudonymous identification. The authors declare that they have no financial interests or conflicts of interest to disclose. This study was conducted as part of a larger research project, which received ethical approval from the Ethics Commission of the Faculty of Humanities at the University of Bern

## 3. Results

In total eight individuals (N female = 2, mean age: 28, SD: 5.7 and N male =6, mean age: 29, SD: 10.2) participants took part in this study. All together, they completed eleven nights at the Institute for Sports Science of the University of Bern. During these eleven nights, a total of six lucid dreams were reported by four participants, verified by eye-movements and/or external rating of the dream report.

Participants reported their dream recall frequency over the past few months. Two recalled dreams almost every morning, three several times a week, two about once a week, and one two to three times per month. One participant frequently experienced lucid dreams (three to five times per week), two occasionally (one to three times per month), and five rarely (less than once per month).

All participants understood the concept of juggling. Juggling proficiency was assessed on a 1–5 scale (1= minimal, 5 = exceptional). One participant rated their skills at level 5, one at level 3, three at level 2, and three at level 1. Five participants could juggle with three balls. Confidence ratings in juggling closely aligned with skill ratings, with only two participants reporting slightly higher confidence than their assessed proficiency.

The following results will be organized in four sections:

- The first section contains non-lucid and lucid dream data. All participants were divided into two groups; the lucid dream group (LD-group) and the non-lucid dream group (NLD-group), based on the experience of a LD in the lab or not.
- In the second section, the LD-group is further explored and split into two groups: high self-reported lucid dream control (LDC-group) and low or no self-reported lucid dream control (NLDC-group). It provides detailed dream reports and further qualitative data about dream control collected from the questionnaire.
- The third and last section focuses on the two LDC-group case reports.

### 3.1 Lucid Dreaming

#### 3.1.1 Induction techniques

In terms of practicing lucid dream techniques, the LD-group showed a more frequent use, with one participant using them frequently (three to five times a week), two participants using them occasionally (one to three times a month), and only one participant using them rarely (less than once a month). In the NLD-group, only one participant uses them occasionally, while the other three participants use them rarely. When asked how they usually become aware that they are dreaming, participants from the LD-group reported heightened awareness through dream signs or anomalies, intentional setting before sleep, identification of incongruities within dreams, and spontaneous realizations. In contrast, participants from the NLD-group cited awareness pathways such as reality testing, spontaneous realization, intuitive awareness, and recurrent reality testing. These results are presented in table 1.

**Table 1.** Induction techniques, the number of participants using this technique, and a rating on how effective they were in brackets.

| Induction technique | LD-group N (rating) | NLD-group N (rating) |
| --- | --- | --- |
| Reality testing | 3 (2) | 2 (5, 5) |
| Noticing the unreal nature of the dream while Dreaming (DILD) | 2 (4) | 2 (5) |
| Intention Setting Before Sleep | 2 (2) | 1 (2) |
| Wake Back to Bed (WBTB) | 3 (3) | 2 (4, 4) |
| Setting an Intention to Remember to Recognize when Dreaming (MILD) | 0 | 1 (4) |
| Maintaining Awareness While Transitioning from Wakefulness to Sleep (WILD) | 2 (3) | 1 (3) |
| Focusing on Sensory Stimuli and Repeatedly Shifting Attention (SSILD) | 1 | 0 |

#### 3.1.2 Reality testing

Every participant mentioned practicing some kind of reality testing to ensure they were in a dream. Searching for abnormal things was the most frequently mentioned technique, with three participants from the LD-group and two participants from the NLD-group rating its effectiveness between 2 and 4. Two participants from the LD-group and one participant from the NLD-group mentioned looking at their hand to check reality, while one participant from the NLD-group tries switching lights on and off. Additionally, one participant from the LD-group uses a talisman for reality testing while dreaming, and one participant from the NLD-group mentioned attempting to fly. Participants were questioned on their use of reality testing during waking life for lucid dream induction and the effectiveness of this technique. Findings revealed that two participants from the LD-group and one participant from the NLD-group engaged in reality testing during waking life.

#### 3.1.3 Dream Journal

Further, participants were asked if they keep a dream journal, how often they use it, and how effective it has been for them. Three out of four participants in the LD-group keep a dream journal and use it between two to three times or less than once per month. The reported effectiveness ranged from 2 to 4. In the NLD-group, only two participants reported using a dream journal, with similar usage frequency (two to three times or less than once per month), and they rated its effectiveness in improving their lucid dreaming skills at 3 and 4.

#### 3.1.4 Dream Goals

Participants provided insights into the establishment of specific goals or intentions for their dreams, with two participants from the LD-group and one participant from the NLD-group confirming goal setting. The success evaluation, rated on a scale from 1 to 5, resulted in varying responses: the LD-group reported ratings of 4 and 2, while the participant from the NLD-group gave a rating of 3.

#### 3.1.5 Motivation and Persistence

For the assessment of motivational persistence, participants rated different statements on a scale from 1 (indicating “very low degree”) to 5 (indicating “very high degree”). The statements were categorized into Long-term Purpose Pursuing, Recurrence of Unattained Purposes, and Current Purpose Pursuing:

- Long-term Purpose Pursuing: The LD-group provided a mean rating of 4.25, indicating strong agreement, whereas the NLD-group had a lower mean rating of 3.25, suggesting a comparatively lower agreement with statements in this category.
- Recurrence of Unattained Purposes: The LD-group exhibited lower agreement, with participants giving individual ratings averaging 2 and a group mean of 2.5. In contrast, the NLD-group had a higher mean rating of 4, indicating stronger agreement with statements related to the recurrence of unattained purposes.
- Current Purpose Pursuing: The LD-group displayed a higher mean rating of 4, suggesting a generally strong agreement. The NLD-group had a slightly lower mean rating of 3.75, indicating a moderate level of agreement with statements related to the pursuit of current purposes.

#### 3.1.6 Self-Efficacy

The participants’ self-efficacy in relation to their juggling skills was recorded, both in lucid dreams and in the waking state. Participants rated their confidence level on a scale from 0 to 100, with 0 representing “the inability to perform the task” and 100 representing “absolute certainty to perform the task.”

##### 3.1.6.1 Juggling in the waking state

The LD-group exhibited a diverse range of confidence levels when considering juggling in the waking state. Two participants displayed contrasting levels of confidence, with one rating at 50 and the other at 100. The remaining two participants expressed lower confidence, both indicating scores of 20.

In the NLD-group confidence levels for juggling in the waking state were more evenly distributed. While two participants showed high confidence with ratings of 80 and 75, the other two participants exhibited lower confidence levels, with scores of 10 and 50.

##### 3.1.6.2 Juggling in a lucid dream

In the LD-group, half of the participants (two individuals) expressed high confidence, both rating their certainty at 100. This indicates a strong belief within these individuals that they can successfully execute juggling while in a lucid dream. The other two participants exhibited moderate confidence, recording values of 40 and 60.

In the NLD-group, participants demonstrated varied levels of confidence. While one participant expressed relatively high certainty with a score of 70, the remaining three participants displayed lower levels of confidence, with scores ranging between 30 and 50. This diversity in responses suggests a spectrum of medium beliefs within the non-lucid dreaming group regarding their ability to juggle in a lucid dream.

#### 3.1.7 Personal beliefs and attitudes

The examination of personal beliefs and attitudes among participants yielded valuable insights into their perspectives on lucid dreaming and its potential impacts.

All participants expressed a belief in their capacity to achieve lucidity in their dreams, as well as in their capacity to juggle within a lucid dream. Six participants believe that lucid dreaming can influence their waking experiences. Two participants from the NLD-group expressed skepticism regarding the potential impact of lucid dreaming on their waking lives.

These findings underscore the participants’ positive attitudes and strong beliefs regarding their ability to attain lucidity, the potential impact of lucid dreaming on waking life, and their perceived capabilities, including the skill of juggling, within the context of lucid dreams.

#### 3.1.8 Stress levels

For the assessment of stress levels among participants, the perceived stress scale was used, wherein participants had to answer ten different questions regarding perceived stress and rate them on a scale from 1 meaning “never” to 5 meaning “very often”. The results revealed variations in perceived stress, categorized into low, moderate, and high stress levels based on the scores obtained. In the following Figure 1, the individual assessed stress levels are shown. The colored background representing the categories of stress level with the withe part being low stress and the orange part being a moderate stress level.

**Figure 1:**
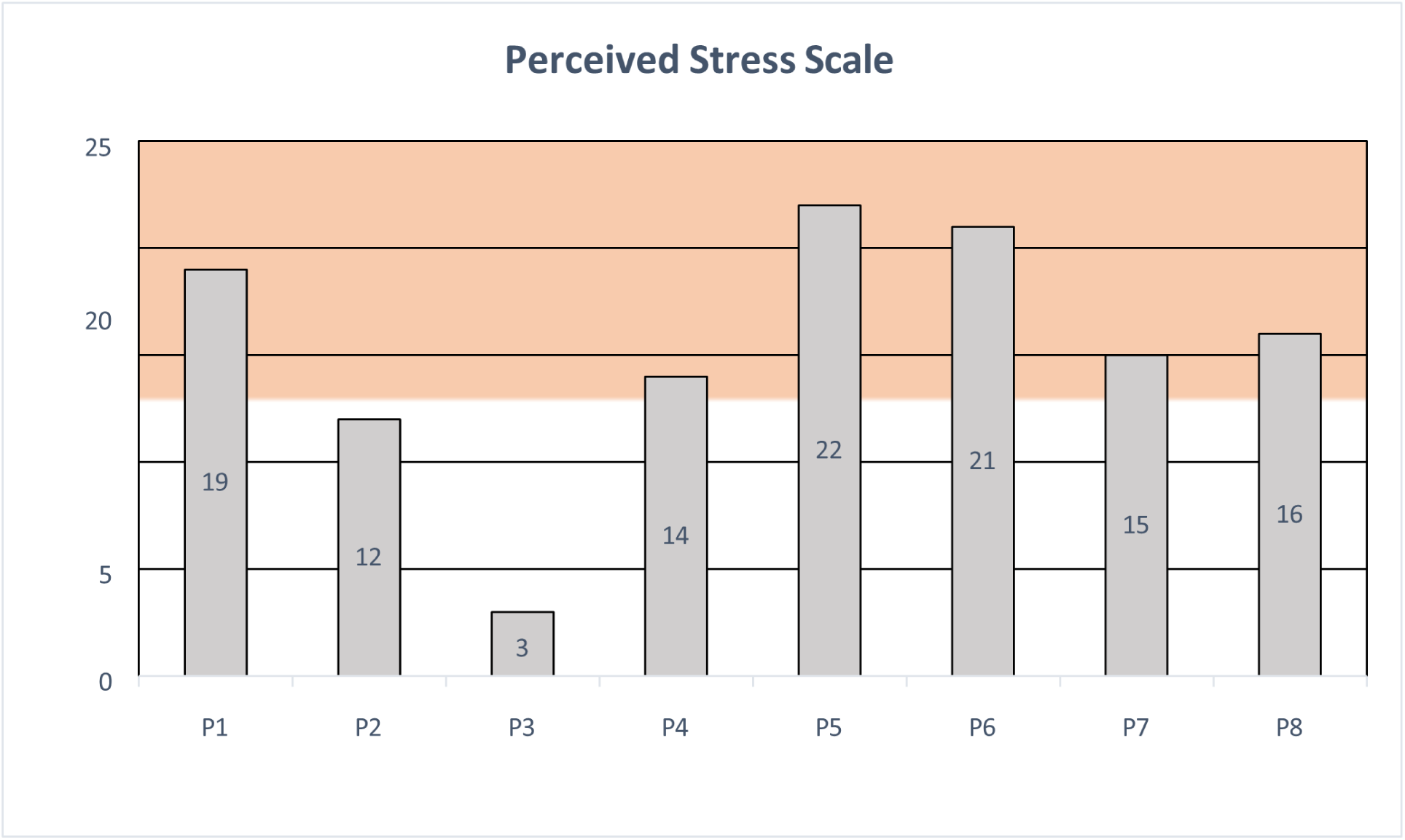
Perceived Stress Scale (PSS) with low stress level in white and moderate stress level in orange.

Low Perceived Stress (0-13): Two participants from the LD-group had scorings under 13 which suggesting that they experienced minimal stress based on the provided scale.

Moderate Perceived Stress (14-26): Six participants fell into the category of moderate stress. Two participants from the LD-group and all four participants from the NLD-group demonstrated stress scores within this range of moderate perceived levels of stress.

High Perceived Stress (27-40): None of the participants reported high perceived stress levels.

The mean values between the two groups revealed a difference, with the LD-group exhibiting a lower value (mean=12, S.D.=6.7) compared to the NLD-group (mean=18.5, S.D.=3.5), whereas the standard deviation is much lower than in the LD-group. In relation to norms established by Cohen and Janicki-Deverts (Cohen & Janicki-Deverts, 2012), the average perceived stress level in the LD-group was more than 4 points below their study’s average (16.78 for under 25-year- olds and 17.46 for 25–34-year-olds). Conversely, the perceived stress level in the NLD-group, with a value of 18.5, exceeded the mean reported by Cohen and Janicki-Deverts (17.46 for 25–34-year-olds and 16.94 for 45–54-year-olds).

#### 3.1.9 Mindfulness

In figure 2, the mindfulness levels are presented with a colored separation between the different categories of mindfulness being D=Describing, OBS=Observing, AA=Acting with Awareness, NJ=Non-judging of inner experience and NR=Nonreactivity to inner experience. One participant from the LD-group expressed low levels of Mindfulness scoring of 26, whilst the other seven participants all scored between 32 and 38 out of 50.

**Figure 2.**
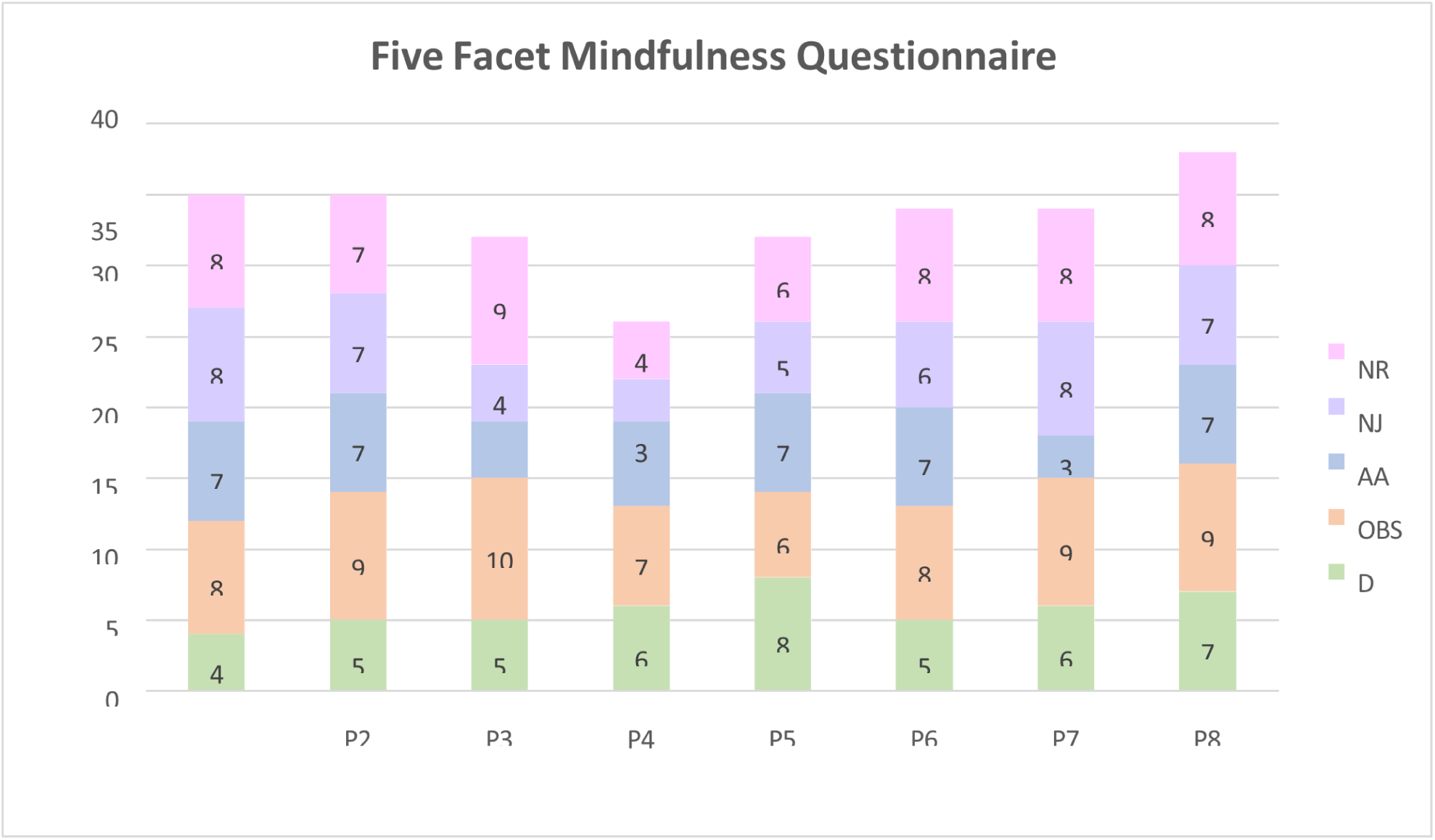
The Five Facet Mindfulness Questionnaire containing every answer, for each Item (each item has 2 Questions scaling from 1-5, max 10 points per Item). Total maximum per person would be 50. D=Describing, OBS=Observing, AA=Acting with Awareness, NJ=Nonjudging of inner experience, NR=Nonreactivity to inner experience.

In the assessment of mindfulness practices, the participants exhibited diverse engagement patterns across various exercises:

- Meditation: One participant from the LD-group meditates frequently (3-5 times per week), Two rarely (less than once per month). From the NLD-group, three occasionally (1-3 times per month)
- Yoga: Two participants from each group practice yoga occasionally (1-3 times per month), and one person from the LD-group practice it rarely (less than once per month).
- Hypnosis: One participant from each group use hypnosis only rarely (less than once per month).

Two participants from the NLD-group reported practicing other mindfulness-related activities such as reading books and singing.

### 3.2 Dream control

In this subsection, only the findings from the participants who experienced a lucid dream within the laboratory setting are presented. A comparison is made between participants demonstrating low or no dream control (NLDC-group, N=2) and those demonstrating higher dream control (LDC-group, N=2) based on their in-lab lucidity experience. Group classification was determined based on the self-reported control ratings in Table 4, where participants rated their level of dream control across different aspects. Those with higher ratings (predominantly 4 and 5) were assigned to the LDC-group, while those with lower ratings (primarily 1 and 2) were placed in the NLDC-group. The total number of participants contributing to this section is N=4.

#### 3.2.1 Lucid dreaming behavior

In the overall assessment of how participants typically behave when they become lucid, a variety of responses were collected. All participants reported having previously attempted to control or manipulate events in their dreams.

- NLDC-group: changing dream scenery or environment (N=2), interacting with dream characters (N=1), manipulating objects or events (N=0), flying or levitation (N=1), and passive observation (N=2).
- LDC-group: changing dream scenery or environment (N=1), interacting with dream characters One, manipulating objects or events (N=1), flying or levitation (N=2), and passive observation (N=2).

#### 3.2.2 Controlling different aspects in a lucid dream

Table 2 presents ratings of the ability to control different aspects of dreams, differentiating between the LDC-group and NLDC-group. Participants were asked to rate their dream control on a scale of 1 meaning “low control” to 5 meaning “high control”. The table shows the average ratings from each group (with the individual ratings in brackets) across various dream control items.

**Table 2.** The subjective rating of the ability to control different aspects of lucid dreams for the LDC and NLDC group. Group average ratings and individual ratings in brackets.

| **'Do you have the ability to control... in your lucid dreams?'** | **LDC-group** | **NLDC-group** |
| --- | --- | --- |
| ...own dream body | 4 (4, 4) | 3 (4, 2) |
| ...other dream characters' bodies | 2,5 (3, 2) | 1 (1, 1) |
| ...own dream body movement | 3,5 (4, 3) | 4 (5, 3) |
| ...own actions | 4 (4, 4) | 3,5 (5, 2) |

#### 3.2.3 Challenges in controlling aspects of lucid dreams

Participants were asked about the challenges and limitations they have experienced while lucid dreaming. Both participants in the LDC-group reported encountering difficulties across nearly all aspects of their lucid dreams, including maintaining lucidity, remembering goals, allowing supernatural events to unfold, staying focused, imagining supernatural actions, and, in some instances, distinguishing dreams from reality. Similarly, both participants in the NLDC-group experienced challenges in maintaining lucidity. However, neither of them reported difficulties in remembering goals or imagining supernatural actions.

Further insights were gained by looking at specific challenges in manipulating different dream elements, showing differences between participants with higher Lucid Dream Control (LDC) and those with low Lucid Dream Control (NLDC).

- LDC-group: Participants in this group generally reported greater challenges in situations where they exerted more influence over the dream. When attempting unfamiliar actions, such as performing a new movement or engaging in supernatural actions, the difficulty increased. While changing environments was possible, it required either a vivid mental image of the location or prior familiarity with the place. In terms of interactions with dream characters, participants only influenced them when necessary. Both participants found it nearly impossible to manipulate the perception of time within their dreams.
- NLDC-group: Participants in this group reported some challenges but rarely influenced anything beyond their own body movements, actions, and minor object manipulations. When engaging in supernatural actions, they noted that gravity became unpredictable or unstable. They generally avoided directly controlling other dream characters and instead influenced them indirectly, primarily through thoughts. Altering the dream environment was considered difficult and only possible when pre-existing mental images or conceptswere available. The perception of time was described as relative and was rarely a focus of their lucid dreaming experiences.

#### 3.2.4 Achievements in lucid dreams

Table 3 provides data into the lucid dream achievements reported by participants. Whereas for the NLDC-group, most actions were achieved by neither or one participant, for the LDC-group more than half of the actions have already been achieved for both participants. Additionally, P2 from the LDC-group reporting an additional achievement of being able to transform into animals.

**Table 3.** Achievements in Lucid Dreams reported by the two groups: LDC and NLDC.

| What are challenges you already managed to do in a lucid dream? | LDC-group (N=2) | NLDC-group (N=2) |
| --- | --- | --- |
| Communicate with dream characters | 2 | 2 |
| Deliberately shape dream environment | 1 | 1 |
| Flying with full control | 2 | 2 |
| Make day turn to night | 2 | 1 |
| Going through walls or objects | 1 | 0 |
| Going through dream characters | 1 | 1 |
| Eat food | 2 | 1 |

Participants reported their lucid dream experience as accurately and extensively as possible followed by answering specific questions regarding lucidity, dream control and other dream content. The dream reports for each participant can be found in the Appendix C.

#### 3.2.5 Awareness

The participants from the LDC-group reported that they mostly get lucid when abnormal things like quick changes of scenery or too many people who should not be in a certain place get detected. “In the beginning I always thought all dreams were real until I realized the difference, that can’t be true.” One of the participants from the LDC-group reported to make a reality check by looking at his right hand and check if something was off. “Then I looked again more closely, and the ring finger or index finger was missing. I just had four fingers. Then I noticed it.” The NLDC-group reported normally reaching lucidity when particularly complicated or abnormal things happen. One of the participants reported to hear voices calling them. “It’s always like as if someone would knock and say: “hey, you’re actually dreaming”.

#### 3.2.6 Eye-signal

Both participants from the LDC-group reported consciously giving the LRLR-eye signal several times within a lucid dream. One participant of the NLDC-group tried to move their eyes but didn’t know if it worked. The second participant from this group reported to not have performed any LRLR eye signal consciously.

#### 3.2.7 Control over dream content

One participant from the LDC-group reported having control over the dream only for a brief period while lucid. During this time, they immediately gave the LRLR eye signal and began juggling. The other participant described experiencing sequences where they could consciously interact with other dream characters and deliberately move away from them. However, they also noted alternating between making intentional choices and passively following the unfolding dream events: “I had to choose a path and just give direction but effectively what I do, I sometimes backed off and left what was coming.” Both participants from the NLDC-group reported no control over the dream content.

#### 3.2.8 Juggling attempt

The participants with higher dream control in their lucid dreams both attempted to juggle. One participant lacked juggling balls and simply mimicked the arm movements without physical objects. The other participant initially tried juggling with standard juggling balls but was only able to control one or two at a time. Later in the dream, they discovered oranges and successfully juggled three of them.

For the self-assessment of juggling skills, only one participant from the LDC-group was able to provide a rating on a scale from 1 (no skills) to 5 (excellent skills). They rated their ability with juggling balls as 2 and their performance with oranges as 4 out of 5. The other participant, having only performed the arm movements without objects, was unable to rate their skills.

Among the NLDC-group, one participant considered juggling but did not follow through. The other participant reported that the idea of juggling never occurred to them and instead waited for external guidance on what to do next.

#### 3.2.9 Stability

In the LDC-group, one participant initially experienced an unstable dream. However, when a dream character (the researcher), who was relevant to the setting, appeared, the dream became more stable. The other participant reported having a stable dream until a dream character (the researcher) covered their eyes, gradually leading to awakening. In the NLDC-group, one participant described their dream as highly stable but with low awareness. The other participant reported fluctuating levels of both stability and awareness throughout the dream.

#### 3.2.10 Intention or goal

All participants in the LD-group set a specific goal before entering their lucid dream. Both participants in the LDC-group had a clear intention for their dream and were able to achieve their goals to some extent.

In the NLDC-group, one participant reported having only a single goal—to move their eyes—but, once in the dream, they no longer remembered the reason for doing so. The other participant intended to juggle but was interrupted before they could follow through.

#### 3.2.11 Other dream characters

All participants reported to interact with dream characters and objects as if they were real. One participant from the LDC-group even described engaging in nonverbal communication, relying solely on body language: “I started talking to her. There was communication, but not verbal—rather through actions and eye signals. Body language.” A participant from the NLDC-group recalled a specific conversation in which they argued with a group of people: “I wanted to go to sleep, but they wanted to stay. I said, ‘No, I want to go to sleep!’”

#### 3.2.12 Emotions

Both groups reported experiencing a mix of emotions during their lucid dreams. Participants in the LDC-group described feelings of nervousness, excitement, fear, hope, and a combination of nervousness and joy. Participants in the NLDC-group reported a broader range of emotions, including love, closeness, disappointment, guilt, a mix of emotionlessness with anger, feelings of unfairness, seeking sympathy, disappointment, anger, and positive emotions.

#### 3.2.13 Time

Regarding the perceived duration of the dream, both groups reported experiencing relatively normal time progression. However, one participant from the LDC-group described a significant contrast between different dreams—one felt as if it lasted for several days, while another lasted only two to three minutes. They explained: “The perception was like in real life, but certain scenes went faster because you are fully in the conversation, and sometimes it felt much slower.” The other participant in the LDC-group estimated their dream lasted only about half a minute. In the NLDC-group, one participant reported their dream lasting two to three hours, while the other did not specify a particular duration.

#### 3.2.14 Colors

In terms of color, one participant from the LDC-group reported minimal colors and lighter, beige, eggshell colors, whereas the dreams from the NLDC-group were reported as ‘colorful’ and ‘very colorful’.

#### 3.2.15 Physics

One participant from the LDC-group reported experiencing a loss of gravity: “The whole room shook, and things suddenly flew around. It became quite intense.” In another dream, they again noticed irregularities in the laws of physics: “I realized that the physics were not right at all.” The other LDC-group participant, however, reported that everything in their dream followed normal physical rules.

In the NLDC-group, one participant described unintentionally breaking the laws of physics by floating in the air and rotating while meditating. The other participant reported experiencing normal physics throughout their dream.

#### 3.2.16 Dream Control

At the end of each dream report, participants were asked to rate their level of control over various aspects of their dream. They evaluated their control on a scale from 1 (“no control”) to 5 (“a high level of control”). Table 4 presents the individual ratings from both the LDC-group and the NLDC-group. A clear distinction emerged between the two groups: participants in the LDC-group consistently reported higher levels of control, with ratings predominantly at 4 and 5, while those in the NLDC-group mostly rated their control as low, with scores of 1 and 2.

**Table 4.**
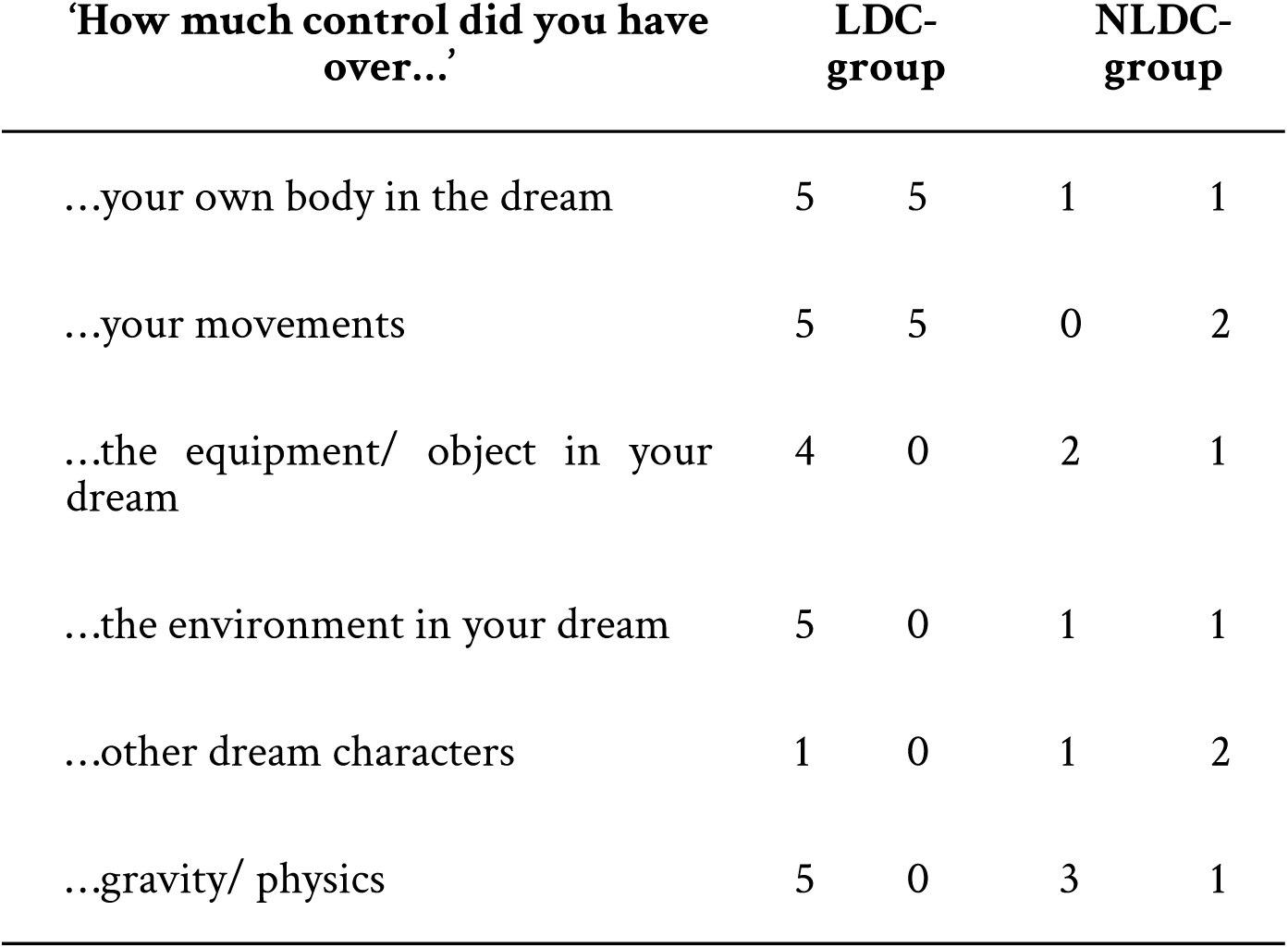
Lucid dream control scaling by the individual participant, separated by group.

### 3.3 Juggling case studies

This section provides a detailed analysis of the two participants in the LDC group, who experienced high dream control and attempted juggling during their lucid dreams.

#### 3.3.1 Participant 1 (P1)

Participant 1 demonstrated extensive experience with lucid dreaming, reporting frequent occurrences several times per week and a strong ability to recall dreams nearly every night. He described typically becoming lucid by recognizing anomalies within his dream. To induce lucidity intentionally, he follows a wake-back-to-bed (WBTB) strategy, setting an alarm at 6:00 AM and pressing snooze every nine minutes until he transitions into a lucid dream. Throughout the study, P1 participated in three sleep lab sessions. Across all three nights, P1 consistently experienced lucid dreams, each validated by the performance of a LRLR eye signal during REM sleep, except for the second night due to technical issues with the PSG recording.

##### 3.3.1.1 2.6.1.1 P1 - Night 1

On the first night, P1 fell asleep without difficulty despite the PSG setup. After approximately five and a half hours of sleep, he was awakened for the WBTB procedure. About 35 minutes after returning to sleep, a REM phase was detected, accompanied by a potential eye signal. Another potential LRLR eye signal was identified at 5:45 AM. At 5:55 AM, P1 reported his dream and confirmed his intent to perform the eye signal. The dream took place in a setting reminiscent of Italy or Spain, involving a secret mission on sleep research, hidden from the mafia. P1 described the dream as highly vivid, including a dream within a dream, featuring decisions and conversations with dream characters. Despite attempting to signal lucidity, he became more awake, but eventually re-entered the dream state.

> “You came in, woke me up, but it was in the dream of the dream.”
>
> “To come back to the lab, I had to choose a path and just give direction but effectively what I do, I sometimes backed off and left what was coming.”
>
> “I realized I was in a dream and told the kid to wait a minute. I moved aside to make the sign quickly. Then I tried a few times and noticed that I was getting a little more awake and went back to dreaming.”

P1’s dream control was primarily centered on influencing decisions and shaping the dream environment, rather than exerting direct control over his own movements or interactions with dream characters. He rated his ability to manipulate both the dream environment and influence gravity as moderate (3 out of 5). However, he reported a lower level of control (1 out of 5) when it came to his own body movements and interactions with dream characters.

##### 3.3.1.2 P1 – Night 2

On the second experimental night, P1 had no difficulty falling asleep. Approximately five and a half hours after sleep onset, he was awakened for the WBTB procedure. Due to technical difficulties, the sleep architecture and potential LRLR eye movements could not be verified. P1 reported experiencing multiple lucid dreams during this night. He became lucid upon noticing an unusual number of people in the dream, which is a common cue for him to recognize lucidity. However, he described his lucid states as short-lived. He also recalled attempting to provide the eye signal and actively setting further goals within the dream.

> “As I was aware that I was lucid, I consciously set goals, for example juggling.”

He wanted to juggle, but after he found juggling balls, he reported that either the dream didn’t want him to juggle, or he was too nervous to do it. He only held the balls in his hands:

> “I first tried to give the sign, and then I wanted to juggle, but I couldn’t because I didn’t have balls and then I had to go looking for balls. Once I had balls, I don’t know if I was nervous, or the dream world didn’t want me to juggle. Then I actually just had the balls in my hand.”

After that, he reported to have woken up, but later realized that this was also a dream. He dreamt of waking up and falling off the bed repeatedly until the researcher came and picked him off the floor. He then fell back asleep, and as he woke up, the researcher was there again to give him oranges because he was hungry. Then he showed his juggling skills with the oranges:

> “I thought I had fallen off the bed, that wasn’t true at all, because then you came and picked me up off the floor again, but then I thought that can’t be true, and woke up and as I fell asleep again, I dove back into a dream and that’s when you were there again. Because I was hungry, you had oranges and then I showed you how I could juggle.”

He reported successfully juggling and exerting some control over gravity, though it felt chaotic. Initially, he tossed a single ball back and forth between his hands, and upon realizing this worked, he attempted to juggle three balls in a circular motion. As this also proved successful, he moved on to controlling the balls with his mind, causing them to move back and forth in a snake-like pattern. He noted that he could alter the weight of the juggling balls depending on the speed at which he threw them. However, upon realizing that this defied real-world physics, he lost control, and the balls dropped to the floor. As soon as he became aware that his actions were illogical, his ability to manipulate the balls stopped.

> “In the beginning the gravity was crazy.”
>
> “I threw the ball normally back and forth in two hands and then I tried, because it was easy, three balls in a circle one after the other. Because that also worked, I threw the balls back and forth with my mind snake-like.”
>
> “It did not make sense afterward, and as soon as I thought it did not make sense, I could control it afterward in the dream. Other things happened around me because it did not work afterward.”

He also reported that sometimes, if he wanted to do something which the dream would not allow, he had to improvise:

> “Sometimes I was aware that I was dreaming, but my body didn’t want to. It was more influenced by the dream world, but I was aware, I have the goal now, but until I reached the goal, it took a little bit.”

He reported having a focused view while the periphery was blurry, so it was easy to focus on the things he was doing.

> “There was also the dream world focused into it. I didn’t have to look at the whole world anymore and then there’s more control.”

He reported excellent juggling skills with the oranges, rating his performance at 4 or even 5 out of 5. In contrast, he assessed his juggling ability with traditional juggling balls as significantly lower, giving it a 2 out of 5. Regarding dream control, he rated his ability to manipulate both the equipment and gravity at 4 out of 5. He reported moderate control (3 out of 5) over his own body, movements, and the dream environment. The least control was observed in his interactions with other dream characters, which he rated at 2 out of 5.

##### 3.3.1.3 P1 – Night 3

On the third and final night, a REM phase was detected five and a half hours after sleep onset, but no eye signal was observed, prompting the WBTB procedure. After a 30-minute waking period, P1 went back to sleep. One and a half hours later (at 6:30 AM), another REM phase was detected. At 7:30 AM, P1 woke up and reported having a short lucid dream that occurred even before the WBTB procedure. In this brief dream, he performed the eye signal, although it could not be precisely identified later in the PSG recordings. He described creating a scene in which he was drinking coffee with a friend.

> “I created a scene. Afterwards I had no control, there was not more than one scene either.”
>
> “It was there, in a room, a table. The scene with the coffee drinking and the one colleague who was standing there.”
>
> “I started talking to her (…) it was with a lot of interaction, there has been communication, but not verbal rather with actions and eye signs. Body language.”

He reported feeling nervous, and described the scene as brief, lasting less than two or three minutes. He also noted that the dream appeared in light colors.

> “More white, beige, eggshell color. The people, me and her, I didn’t look at me, but she looked like she used to (…) the environment was white because I created that.”

He reported experiencing some difficulties with dream stability as his lucid experience ended. “The more I wanted to change or create, it wasn’t stable afterwards, I woke up right after.”

He reported to have a very high control over almost every aspect asked, with control over the own body, the movements, environment, and gravity rating with the maximum of 5 and control over objects he reported a 4. Because he didn’t want to influence the other dream character, he rated the control over this aspect with a 1.

#### 3.3.2 Participant 2 (P2)

The second participant (P2) spent two nights in the sleep laboratory, achieving a lucid dream on the second night. P2 had no issues falling asleep and experienced multiple REM phases. After approximately three and a half hours post-sleep onset, an LRLR eye signal was observed and detected, as shown in Figure 3.

**Figure 3:**
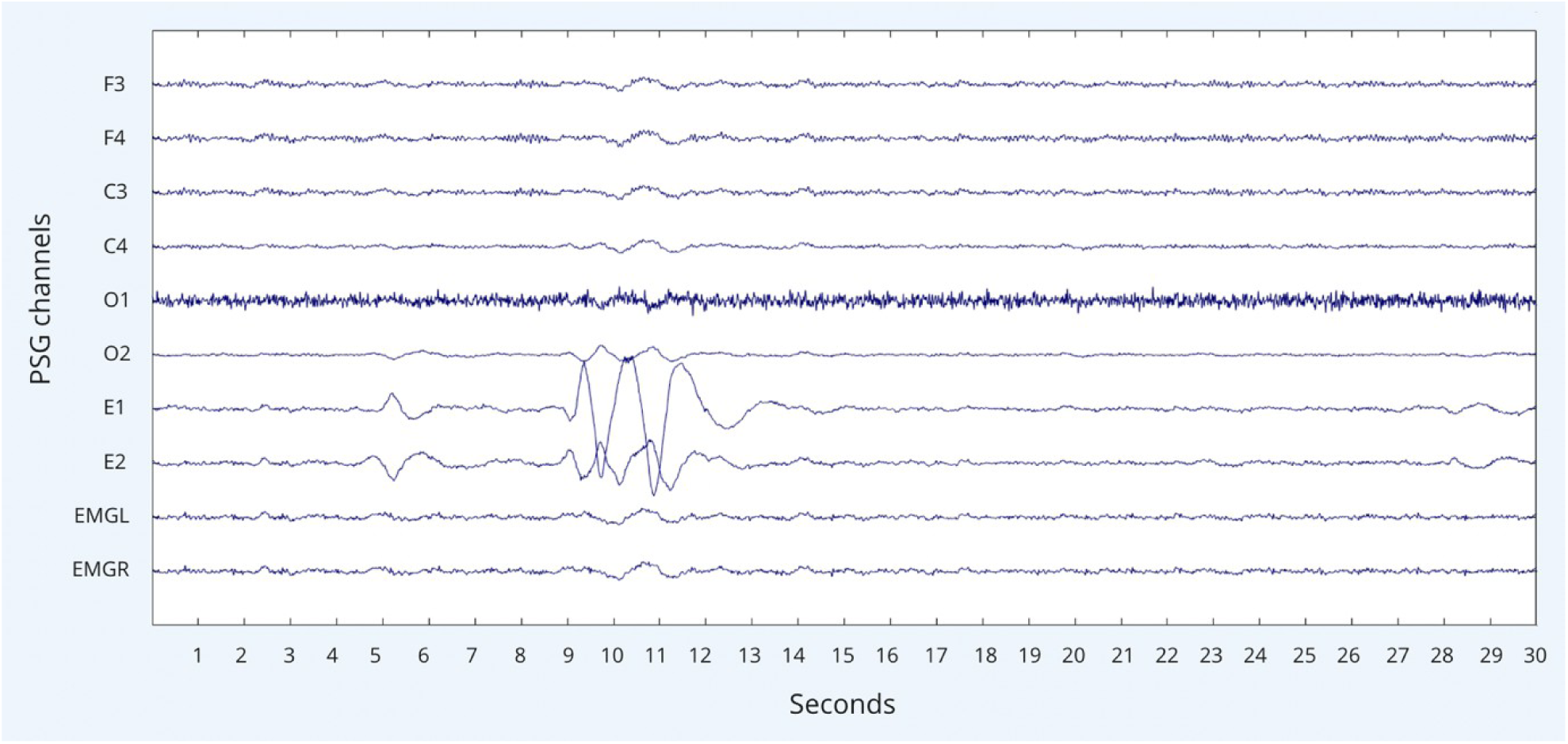
Signal-verified lucid dream (SVLD) in REM sleep of participant 2 – night 2 at 2:22 (with EEGLAB)

Approximately 20 minutes later, a dream report was recorded in which the participant reported giv-ing the eye signal several times.

“Did you receive my signal?”

> “I gave it several times. I had a lucid dream and did the juggling movement.”
>
> “I looked at my hand and I had four fingers. I did the eye movement quickly, left, right, left, right. I did it several times. Then I did the movement right away.”

P2 dreamed of being in the laboratory setting, sitting on the edge of the bed next to the researcher. To verify whether he was dreaming, he checked his hands and became excited upon discovering that he had only four fingers, confirming that he was lucid dreaming:

> “I looked at my right hand, and it didn’t seem like a dream at all. Then I looked again more closely, and the ring finger or index finger was missing.”

During the dream, he attempted juggling movements with his hands, but without any juggling balls. While doing so, he repeatedly performed the LRLR eye signal. However, his focus was interrupted by a dream character:

> “I did the eye movement quickly—left, right, left, right. I did it several times. Then I did the (juggling) movement right away. While I was doing it, I kept doing LRLR in between. You (the researcher) turned to me and put your hands in front of my eyes. You were trying to stop me from doing something.”

Shortly after this encounter, P2 believed he had woken up and spoke to the researcher, who supposedly entered the room and removed his electrodes. However, this turned out to be a false awakening, as the researcher had not actually entered his room. Only after waking up for real did he conduct the dream report:

> “I thought I woke up because you came into the room, and I asked, ‘Hey, did you get it?’ You said, ‘Yes.’ Then we high-fived. But in this case, this was still a dream too. I even ripped the electrodes off here—funny.”

P2 recognized that his dream was short, which motivated him to attempt juggling quickly. Even though he had no balls, he initiated the juggling movement, hoping that balls might appear—but they did not:

> “I thought that maybe the balls would come, but they didn’t. I thought, ‘Since I can do it myself, it won’t be much different.’ So, I just did the movement.”

He reported that his juggling skills were the same as in waking life, as did his body movements and the effects of gravity. Everything felt normal, and he had full control over his body. However, he did not attempt to control other dream characters, objects, or gravity, so he could not assess his level of control over these aspects.

P2 described experiencing full clarity during the dream and feeling joy upon realizing he was lucid. However, he quickly refocused to carry out his intention to juggle without exerting additional influence on the dream. He reported that the entire lucid episode seemed to last approximately 30 seconds.

## 4. Discussion

The ability to manipulate dream content deliberately in a lucid dream has significant implications for skill improvement, problem-solving, and mental well-being. While much of the research on lucid dreaming has focused on induction techniques, dream control remains a crucial factor that determines the efficacy of LD-based interventions. This study aimed to examine how different psychological, cognitive, and experiential factors contribute to dream control and motor practice in LD.

It is important to note that as our data is based on a very small group of participants, statistical testing was not possible and thus we cannot make any conclusions. Nonetheless, we will discuss our methods, findings and ideas and hope to boost the conversation on moving towards more in-depth research on the factors potentially contributing to the wide variance found in dream control.

Eight participants were included in the study, but only half became lucid in the lab setting. Participants who successfully entered a lucid dream in the lab (LD-group) slightly differed from those who did not (NLD-group) in terms of beliefs, attitudes, and self-efficacy. Confidence in LD ability was generally high, but members of the NLD-group exhibited more skepticism about lucid dreaming’s impact on waking life. This skepticism may reflect doubts about the stability and controllability of LD experiences, and therefore it’s applicability. Prior research has shown that individuals who view dreams as meaningful and controllable tend to report higher lucid dream frequency and stability (Cernovsky, 1984; Schredl et al., 1996). Moreover, a strong interest in dreams has been linked to increased dream recall (Schredl, Ciric, et al., 2003; Schredl, Wittmann, et al., 2003), reinforcing the importance of cognitive engagement in LD success. This is the reason we included them in the assessment; they might improve not only dream recall and lucid dream frequency, but also control of the dream. This could be tested in a future study.

Six out of eight participants rated their ability to juggle in an LD as higher than in waking life, with the only two exceptions belonging to the NLD-group. This could be explained by the fact that the NLD-group had less experience with LD and thus less confidence. Here, it is probable that a high LD frequency increases a person’s self-efficacy of a certain LD task. The other way around, it remains speculation whether self-efficacy predicts a higher level of dream control. It might align with the selfefficacy theories (Bandura, 2005), suggesting that belief in one’s ability to manipulate dream content plays a central role in achieving control.

Additionally, the LD-group reported lower stress levels than the NLD-group, with their scores falling below the normative range for their age group (Cohen & Janicki-Deverts, 2012). These differences could not be statistically verified, but they might align with previous research linking emotional regulation and stress reduction to greater LD frequency and stability (Doll et al., 2009; Gackenbach & Bosveld, 1991). Potentially, reduced stress may facilitate dream control by minimizing stress-related disruptions that can lead to premature awakening or loss of lucidity.

This section examines the results related to dream control, based on the results from online survey responses and participants’ dream reports. To analyze differences in dream control ability, participants in the LD-group were further categorized into those with low dream control (NLDC-group) and high dream control (LDC-group).

Participants who became lucid in the lab exhibited varying levels of control, with some successfully modifying environments and executing complex tasks, while others struggled with maintaining lucidity or carrying out intended actions. Those in the LDC-group reported a greater ability to manipulate dream content; however, paradoxically, they also encountered more challenges. This may be due to the increased cognitive demands required to modify the dream world while maintaining stability and lucidity. A key difficulty reported by participants was the ability to create novel dream elements. While modifying familiar aspects of the dream (e.g., known environments, people, or objects) was relatively easy, generating entirely new scenarios or unfamiliar objects proved more difficult. Many participants also found that larger-scale changes led to dream instability, making it harder to sustain lucidity. Both groups also reported that manipulating familiar dream elements—such as known environments, people, or objects—was significantly easier than making more abstract or extreme modifications.

To counteract these difficulties, participants frequently adopted a “go with the flow” strategy to stabilize their dreams and prevent premature awakening. Their ability to manipulate the dream environment seemed to be limited by their belief in the plausibility of dream content. While those with higher dream control exhibited greater confidence in their ability to influence the dream, they still struggled with actions that conflicted with pre-existing beliefs about reality. Mallett (2021) highlight the complex relationship between dream bizarreness and lucidity, suggesting that higher dream bizarreness often correlates with stronger lucid dream experiences. However, they also indicate that greater lucidity does not necessarily translate into greater dream control, as we already pointed out in the introduction section, particularly when attempting actions that conflict with waking-world expectations. This is what we see in the reports of the current study, where participants reported greater difficulty executing actions they perceived as “unbelievable” or highly unrealistic. While more conventional motor tasks—such as throwing a ball from one hand to the other—might be more achievable, actions that fundamentally violated physical laws (e.g., moving the balls in mid-air like a snake) seem more difficult. This suggests that pre-existing cognitive schemas about reality persist in the dream state, influencing what is perceived as possible even within lucid dreams.

Mallett et al.’s findings align with this observation, as their study suggests that dreamers’ implicit expectations about reality shape the degree of control they can exert. When an action is perceived as too implausible, the dreamer may encounter resistance or lose control. This highlights a potential cognitive barrier to lucid dream motor training—in order for dreamers to fully use lucid dreams to their potential, they may first need to train themselves to override their pre-existing waking-world assumptions about what is possible. Future research could explore whether gradual exposure to increasingly “impossible” dream tasks (e.g., starting with minor dream manipulations before attempting extreme modifications) could help lucid dreamers expand their range of control and reduce the cognitive resistance associated with bizarre or unrealistic dream actions.

Participants’ experiences align with the findings of Mallett (Mallett et al., 2021) which shows that lucidity exists along a continuum rather than as an all-or-nothing phenomenon and this extends to lucid dream control. Participants in the NLDC-group tended to be passive observers, engaging minimally with their dream environment. In contrast, those in the LDC-group attempted more extensive alterations, often resulting in instability or loss of lucidity. Participants’ previous dream achievements further illustrate these differences in control. Many successfully engaged in conversations, environmental manipulation, altering lighting conditions (e.g., turning day into night), and even flying with full control. While some of these activities are fantastical, they still retain elements of physical experience, which may make them easier to conceptualize and execute within a dream.

### 4.1 Limitations

The study faced inherent constraints due to the rarity of lucid dreaming. Finding eligible participants was challenging and not all individuals possessed the ability to become lucid in the lab limiting the amount of usable data. As a result, the sample size remained too small to draw any meaningful conclusions and making broader generalizations more difficult. In addition, limitations in the questionnaire design must be considered. Not all participants provided complete responses on every question, and particularly those question related to effectiveness or proficiency levels, were left unanswered. This incomplete data limited the ability to conduct a thorough comparison of the effectiveness of lucid dreaming in motor skill improvement. Additionally, the questionnaire included study-specific questions, some of which were newly developed rather than adapted from validated scales. Ideally, these new scales should have undergone validation before implementation to ensure reliability and accuracy. Furthermore, the online format limited participants’ ability to seek clarification if needed, which might have affected the precision and consistency of their responses. Future studies could build on the proposed items by validating them through pilot testing, making them more robust for systematically investigating the cognitive constraints of lucid dream motor practice.

### 4.2 Conclusion

While the data in this study was not tested for statistical significance due to the low number of participants, the methods are a step towards more research on dream control. Our findings suggest that dream control is a skill that may develop over time. Participants with greater LD experience and higher dream control encountered more challenges, possibly because they actively pushed the boundaries of dream manipulation. This raises an important question: Can dream control be systematically improved through training? If a higher frequency of lucid dreaming leads to more exposure to challenges, it may facilitate greater overall dream control. Future research could explore whether structured LD training can enhance dream manipulation skills and whether certain cognitive strategies (e.g., belief modification, visualization, or expectation-setting) can improve the ability to perform complex tasks in lucid dreams. In addition, it should be systematically tested whether the level of dream control is dependent on the cognitive parameters mentioned in this study. By examining how various psychological and cognitive factors influence dream control, we can identify key predictors of dream control success and overcome challenges in executing complex dream tasks for the various LD applications.

## 6. Appendices

## 6.1 Appendix A: Lucid Dream Skill Questionnaire (LUSK) Results

| Participant | CODE | age | gender | LD Frequency | In how many of your lucid dreams... |  |  |  |  |  |  |  |  |  |
| --- | --- | --- | --- | --- | --- | --- | --- | --- | --- | --- | --- | --- | --- | --- |
|  |  |  |  |  | Aware of differences to the waking state (e.g., bizarre incidents or settings)? | Aware that your physical body was asleep | Aware that all dream objects were not real | Thought about different options of what you can do in a dream | Able to keep your awareness for a satisfying period of time | Decided deliberately to observe the dream as a dream | Shape your environment (e.g., change landscapes/surroundings, let persons/characters appear or disappear) | Have full control of your dream body (movements, actions) | Chose deliberately a specific action? | Perform specific actions (e.g., flying, floating, talking with dream characters, perform magic, engaging in sex) |
| P1 | SUCHFE | 25 | male | Several times a week | In half | In none | In none | In all | In a quarter | In half | In half | In half | In half | In half |
| P2 | SAANBE | 25 | male | Two or three times a month | In three quarters | In three quarters | In three quarters | In three quarters | In three quarters | In a quarter | In half | In half | In three quarters | In three quarters |
| P3 | COMAAV | 24 | male | About once a week | In three quarters | In half | In a quarter | In none | In half | In all | In a quarter | In three quarters | In all | In all |
| P4 | LUANDER | 24 | female | About two to four times a year | In half | In half | In a quarter | In half | In a quarter | In a quarter | In a quarter | In a quarter | In half | In half |
| P5 | PAFRGI | 25 | male | Less than once a year | In three quarters | In three quarters | In all | In half | In a quarter | In none | In three quarters | In all | In all | In all |
| P6 | MAHAPE | 50 | male | About two to four times a year | In half | In half | In three quarters | In a quarter | In a quarter | In a quarter | In a quarter | In a quarter | In a quarter | In a quarter |
| P7 | ROPEZI | 26 | male | About two to four times a year | In all | In all | In a quarter | In half | In all | In three quarters | In none | In three quarters | In three quarters | In all |
| P8 | DEAELA | 32 | female | Two or three times a month | In a quarter | In none | In a quarter | In half | In half | In none | In three quarters | In half | In half | In three quarters |

## 6.2 Appendix B: Dream Report Questionnaire

Open Question: What went through your head before you woke up? Specific Questions:

On a scale from 1-5, how clear was your dream?

- Did you become lucid?
- How did you become aware that you were dreaming?
- Could you give the eye signal?
- Did you have control over the dream content?
- Did you try to juggle?
- On a scale of 1-5, how good were your juggling skills?
- With what kind of object or item did you juggle?
- Did you have a clear intention or goal for the dream?
- How vivid was the dream environment?
- Were you able to in interact with dream characters and objects as if they were real?
- Did you experience any negative emotions during the dream?
- How long did the lucid dream last?
- Did you dream in color?
- Were the physics in the dream normal?
- Did time move normal?
- Was the dream stable?

Motor task related questions:

Rate these items from,1-5: How much control did you have over…

- … own body in the dream?
- … your own movements?
- … the equipment/objects in your dream?
- … the environment of your dream?
- … other dream characters?
- … The physics/gravity of the dream?

## 6.3 Appendix C: Translated and Converted Dream Reports

### 6.3.1 Participant 1 - Dream report 1

You woke me up but just before at the very end I realized that I am still in the dream. In this scene I was already once. It was in the sleep lab, but the sleep lab was not here but in Italy, Spain area on a hill. Next to the hill was a house protected by the military. We did experiments, but you still have a father and mother who are with the mafia. He doesn’t want you to do experiments, and that’s why the lab is hidden at a like miniature golf, similar setup with roller coaster hidden in the eighteenth hole, with a giant mill. It was my task to lucid dream and that’s why I was by my bed and fell asleep I almost made it once and the second time I made it in the dream. Then I didn’t even wake up and you came in, woke me up, but it was in the dream of the dream. You woke me up and I woke up. He told me that it was good, you can take a break now, go out for a run, but you have to be careful and because I didn’t have any pants, and you gave me a pair. I had to be inconspicuous. We went out with your father and mother. I wanted to go back to the lab, but I didn’t have a key. You had to be inconspicuous like one, beggar would give the money, you handed it to me and your father noticed the pants because they were so perfectly tailored for him, and he noticed because they are his pants. I imitated certain poses like a beggar and then he said yes go. To go back it was bright because it was like in a battlefield. On a hill like a castle or hut, I had to overcome this first. There had formed some group of villagers. They were not good, sometimes they shot themselves or they could not handle the gun. I kept running on the side towards the sleep lab. Then I could sleep again. Before I got to the sleep lab. I was on the bus to get here; I saw a couple of assistants who were also on the bus, and they knew a better way to get to the sleep lab. He said you must get off earlier and then I got off before and it was a new scenery, Italian vibes. I realized I was in a dream and told the kid to wait a minute. I moved aside to make the sign quickly. It was a very unfamiliar sign to make and continue dreaming, quickly I no longer knew how to do it. Then I tried a few times and noticed that I was getting a little more awake and went back to dreaming. Then I was already in the sleep lab, it went on with the dream within the dream. Before you woke me up, I was in some kind of sleep paralysis? I was in that state, you actually had to wake me up a little bit more effectively. The bus scene was only short, I just gave the signal and let it continue. I got to the sleep lab thinking I was already awake. Once I started dreaming again, I had a little more control. Before the dream of the dream, I dreamed that I was in the lab and not specially equipped because the whole room shook, and things suddenly flew around it became quite intense. You were supposed to wake me up for real, I had no control. In the second dream I had more control, in the communication between you and me I consciously moved away from you a little bit because of your father. To come back to the lab, I had to choose a path and just give direction but effectively what I do, I sometimes backed off and left what was coming.

### 6.3.2 Participant 1 - Dream report 2

167 The first dreams were about juggling, I was in the dream, and I realized that I was in the dream. Then I first tried to give the sign and then I wanted to juggle, but I couldn’t because I didn’t have balls and then I had to go looking for balls once I had balls, I don’t know if I was nervous, or the dream world didn’t want me to juggle. Then I actually just had the balls in my hand. Certain things in the dream I knew were not true and then I thought: ‘aha’. In the beginning I always thought all dreams were real until I realized the difference, that can’t be true. For example, in the dream there were way too many people who are actually not there at that time. Then I thought I was waking up, but in the end, I was in a dream and I had already fallen on the floor a few times, but I didn’t fall on the floor at all, I was always in bed. I juggled with normal juggling balls. In the beginning I only had, even though I was looking for three, I only tried with one or two because I can’t juggle properly. Once I managed, but it was not on the first or second try, but on the third, with oranges. I didn’t have to look for them, I found them because I was hungry. The process of waking up, I actually didn’t wake up from the dream, I just kept dreaming. I thought I had fallen off the bed, that wasn’t true at all, because then you came and picked me up off the floor again, but then I thought that can’t be true, and woke up and as I fell asleep again, I dove back into a dream and that’s when you were there again. Because I was hungry, you had oranges and then I showed you how I could juggle. As soon as I was aware that I was lucid, I had consciously set some goals, for example juggling. But there were always some difficulties that sometimes I was only lucid for a short time. As soon as I wanted to do something that the dream wouldn’t allow, I had to improvise a little. When I was not lucid, it was very clear with certain details. Once I became lucid, the focus zoomed in and on small details and the big hole was no longer there or at least blurred. At the beginning I was nervous, exited, then a certain fear when, for example, I fell off the bed and still couldn’t get up, like I was paralyzed and had to wait for you. But then when you came over. It went on forever. Already several days, two days. For example, in the beginning the gravity was crazy because sometimes when I juggle, I throw the ball normally back and forth in two hands and then I tried, because it was easy, three balls in a circle one after the other. Because that also worked, I threw the balls back and forth with my mind snake-like. That also worked and then I realized that the physics is not right at all, and the balls usually fell to the ground. Weight of the ball I could determine in each case according to the speed when throwing, that it becomes lighter. It did not make sense afterwards and as soon as I thought it did not make sense, I could control it afterwards in the dream. Other things happened around me, because it did not work afterwards. Sometimes it took me a little while to realize it was a dream. It always felt very real until I realized that couldn’t be true. Sometimes I was aware that I was dreaming, but my body didn’t want to. It was more influenced by the dream world, but I was aware, I have the goal now but until I reached the goal it took a little bit. The things I was controlling, balls and such, I was already very focused on that. There was also the dream world focused into it. I didn’t have to look at the whole world anymore and then there’s more control. I looked for a setting and then it happened automatically and I didn’t have to change much afterwards. When I was looking for the balls or trying to juggle I had to set a lot of things in motion and that gave a lot back. The more I changed things, the harder it became to juggle.

### 6.3.3 Participant 1 - Dream report 3

It was a housewarming party of a colleague; we supported her there. I crept back and forth, watched and supported the one colleague to change the lamp. The one who calls her did not show up, then I had to support. The colleague’s name is *colleague*. The people introduced themselves, then I said that I am *Participant 1*. I saw another triplet. Then I could only consciously approach next to that particular scene, it was very fast until I woke up again. Only in that one particular scene. I created a scene, afterwards I had no control, there was not more than one scene either. It was there, in a room, a table, the scene with the coffee drinking and the one colleague who was standing there. The lucid one wasn’t very vivid, it was just the two of us. It was just empty. At the housewarming party it was very lively and very many people, familiar and unfamiliar faces. In the lucid dream, in the beginning there were only two of us and I deliberately created the coffee scene. I started talking to her, but this was very short, the dream, and it was after with a lot of interaction, there has been communication, but not verbal rather with actions and eye signs. Body language. With the other one, there has been a lot of talking. In the beginning I was in a main hall, suddenly *colleague* came out of the ceiling and said: “yes *Participant 1*, can you help me quickly assemble the lamp” and I so “yes”. When I was done, I went outside and there were a lot of people gathered on the sofa. The house was very big, and *colleague*’s mother came and introduced herself to the people. Swords, her brother and they all looked exactly the same from the face. I don’t know her family at all, but I know the mother and they all had the mother’s face, but tall, short, fat, it was funny. The sister introduced triplets she has, but no sister has all the triplets. Meanwhile, I woke up. In the lucid dream with the colleague, I go for a coffee today, I was a little nervous I would say. At the housewarming party I had some nostalgia of meeting people from the past and some joy. The first one had the people, me and her, they didn’t look at me, but she looked like she used to, a little bit colored but the environment was white because I created that. The second one, it was super-duper multicolored. She was not coming down from the ceiling, she was in the ceiling and opening up the ceiling. Before she opened the ceiling, I was alone in that room. Because I heard a noise. Then I was walking back and forth to see where the noise was coming from and then suddenly, she opened the blanket and said: “*Participant 1*, can you quickly” she first asked where the other colleague was, I asked her “probably outside smoking” and that she said, “help me quickly” and I was like “okay”. There the gravity was all right. Overall time would have been normal. But certain scenes went faster in that sense, others were in a normal time frame. The perception of me was like in real life, sometimes things pass faster because you are fully in the conversation and sometimes, I thought: ‘when this is finally over?’ I had the feeling that the short dream when I was lucid felt much slower, that time was not passing like in the dream now. The more I wanted to change or create, it wasn’t stable afterwards.

### 6.3.4 Participant 2 - Dream report 1

It was a summer evening, and we jumped from somewhere into the lake, lucidity was not there. However, two hours after I fell asleep, I had the same scenario as last time. You would have woken me up, only I didn’t do a reality check and didn’t realize it was a dream. Well, before I woke up another Stranger came up to us while we were jumping down, asked “do you guys want this car? It’s still in good condition, I just don’t want it anymore.” We were like “no, no, it’s fine” then he said, “the car is right over there don’t you want to go look at it?” Then a woman who was also with us, “yes I would like to see it too” she had such small tusks between her incisors. Signs would have been there for a reality check, but I’m so full of like “oh car, okay, let’s go take a look”. The car ran well, very well in fact. I was with my brother, my girlfriend and there was someone else, but I don’t know exactly who, then the unknown woman who came later while he (the man with the car) was talking to us.

### 6.3.5 Participant 2 - Dream report 2

Did you receive my signal? I gave it several times. I had a lucid dream and did the juggling movement. You were sitting next to me doing something, I looked at my hand and I had four fingers. I did the eye movement quickly, left right left right. I did it several times. Then I did the movement right away. While I was doing it, I kept doing LRLR in between. You turned to me and put your hands in front of my eyes: “laugh”. You were trying to stop me from doing something. Then it was over. I meant I woke up because you came into the room and I asked, “hey did you get it?” you said “yes”. Then we high fived. In this case, this was still a dream too. I even ripped the electrodes off here, funny. I looked at my right hand and it didn’t seem like a dream at all. Then I looked again more closely, and the ring finger or index finger was missing. I just had four fingers. Then I noticed it. It was only a short section, but there I had control. There I was lucid. I did the eye sign right away, remembered it. Then started juggling. I wanted to start it really fast, not that I find distractions. I think at first that in the course, balls would come but they did not. I thought, but because I can do it myself, it won’t be much different. I thought I’ll just do the movement. It was just the left side, which I perceived because I was sitting down. You were in front of me doing something at the table with the cables. I was looking down. I was sitting to the left, that’s all I perceived. I was simply carrying out the intention, not much else. The moment of surprise that I look at the hand. Joy in the sense of “Yeh” and “okay, okay, I have to do this now”. It felt like I didn’t have much time, and I wanted to do as much as possible in that time. It was not the stability. Like it was going down, that moment when the person turned around and covered my eyes, I knew it was slowly over. I didn’t even think about moving anywhere else. The sequence was not so long. Maybe half a minute. From the first time LRLR to the last LRLR, I was steadily making the juggling motion. It had normal color. It was bright, the wall was white. What you were wearing I don’t know; I think it was a dark shirt. I can’t tell clearly. There wasn’t much gravitational stuff going on either. I just sat there and made arm movements. It didn’t have any balls, unfortunately.

### 6.3.6 Participant 3 - Dream report 1

I was here, I had to get up. I didn’t realize it was a dream; I thought I had to leave, then we talked. I left because I thought I was dreaming and then I went to the desert and dreamed of meditating. Then I tried to do something with my eyes, but I don’t know if I managed it. I didn’t want to influence or try to influence the dream. I just know that afterwards we ate a pineapple that weighed 13kg. I kept trying to move my eyes. I was meditating on a balcony on a desert in a palace. I was turning with my body. I always thought I had to move my eyes. Whether it worked or not, I don’t know. At the very beginning I dreamt that you woke me up and then you said that we have to go now, then I sat. Someone has to take the things off my head. You came back and took the things away then we talked. You told me that you fancy me, I said that I noticed this, and we should kiss once. But it won’t have a future I said and now I have to pack my stuff and go. Then another guy I knew came in. He asked what I was still doing here, I should pack my stuff. Then I was standing here in the kitchen and I still had to wash my stuff. That was the moment when I realized “Yes, you are dreaming, you haven’t left yet, and you are still lying here in bed”. Then the switch took place when I was suddenly meditating in the desert. The beginning with getting up felt very real and towards the end it was unreal. The one in the desert was like a narration, as if you were listening to a famous story from a meditation story. Then it was surreal, it became more and more surreal. It’s always like this, the thought flies through my head as if someone would knock on it and say “hey, you’re actually dreaming”. It’s exactly the same as if someone said “hey, you’re dreaming”. I say it to myself, but I don’t know what impulse makes it happen. When I talked to you, I had feelings of love and closeness, but also some disappointment towards you, it felt weird. I felt guilty towards you, and you said it was all okay, but I didn’t really believe that. Then we were okay, and I left it on the side and was washing up. At the end of the story, I was emotionless. The dream happened, as if by itself and I tried to influence, I didn’t specifically try to do anything. My only goal was to move my eyes, I didn’t know why anymore. Juggling didn’t cross my mind. I was just moving my eyes and then waiting for someone to tell me what to do. When I was meditating, it was already in the desert in the palace, the people, I don’t know if it was me or not, I was like floating and I could turn while floating. Then when I had the thought that I had to move my eyes, I also turned in the meditation position.

### 6.3.7 Participant 4 - Dream report 1

I had the feeling that I was half aware that I was dreaming at one point, because it was so long and complicated. I also dreamed about the laboratory. There were a lot of people here. First, I woke up as a test person, then there were all these people around, I half recognized them, they were all from a school. I wanted to go back to sleep, but the people were doing something, they were having a seminar in this room I was in. Another room opened up, there were still a few playing basketball. “laughed” I actually wanted to go back to sleep and then we all went for a walk together. It was all very realistic. It all made sense. Because everything was in the lab, I didn’t feel like I was dreaming. Then I fell asleep again in my dream and then I realized that I was dreaming but it was too weird, I couldn’t really do anything. I actually went to sleep in a dream and then, while falling asleep, I realized that I was dreaming because I fell asleep strangely and woke up again and then was awake really quickly. I had the intention in my dream but thought I was awake and dreamed about the same situation in my dream. I was disturbed by quite a lot of people in my dream. I was actually trying to juggle. I didn’t get to it. I first dreamed of the laboratory, and everything was very vivid. Now it comes to my mind, there was a little dog. I was petting it and talking to people, it was all very lively. Then we went for a walk, it was all very realistic and very detailed. I talked with quite a lot of people and also with you, you were there too. I discussed with people, with one of them I even, now it comes back to me, I was in the lab, and I wanted to go to sleep and there were people sitting on my bed and I had to shoo them away and the one I had to push away a little bit because he was lying on my bed. My bed was in the middle of the room and there were lots of desks all around, like a seminar room. They wanted to do a session there and I wanted to go to sleep, they wanted to be there, I said “no I want to go to sleep!”. Then we made a compromise that we would all go for a walk. When I wanted to go to sleep, I was angry at the people for all of them being in my room. Then I argued with them, I thought it was unfair, I tried to show them that it is very important to me that I can go back to sleep now, they should wait a few more hours. I was looking for sympathy, I was a little disappointed and angry at people. Afterwards I also had positive emotions when they were more empathetic after all, and we looked for a solution. When I woke up in my dream, I felt like it was time for WBTB and I was awake for the whole morning, it felt like it was 2-3 hours. It was very stable, I was not directly aware that it was a dream, so I was just dreaming, and I experienced everything strongly. It was very stable.

### 6.3.8 Participant 4 - Dream report 2

I was just on a ski lift, but it was summer. It was a little weird, we were walking around this place. Because it’s a ski resort, it had ski lifts, and they were running, and we were delivering beer. I was with people who I didn’t know earlier, who I met along the way. It was like a tornado Selzer but a different brand, a new beer. We delivered this beer in this place. Afterwards we were there fooling around, and it had a ski lift, which we were now sitting on. It was very vivid; I perceived many details. It was in nature, when I looked around, everything was very realistic, I saw a little of this place. When I focused on it, I don’t know if it was really this place but the way I imagined it, I saw mountains and recognized the ski lift, everything made a lot of sense. I interacted with people a lot, at nonsense and brought beer. Now it comes back to my mind, we were mixing a syrup, and it was about what color it was. We were making fun of certain people’s color perception. For me it was pink, someone said it was green, one man said it was white. It was funny, I interacted with a lot of people. We were doing some nonsense and had quite fun together, teasing each other and generally good mood, my emotions were also positive.

### 6.3.9 Participant 5 - Dream report 1

I was at the beginning of a dream, but I didn’t do anything yet. I incorporated the laboratory situation into the dream. I was having a hard time falling back asleep. I believe it was true. I don’t think I simply imagined this. I believe I experienced a prolonged period of drowsiness. I thought I had already been woken up. I imagined this situation where there was another researcher, in addition to *researcher 1*, undefined. they had woken me up and no one came to “assist” me in the process of induction; so, the Wake-up-back-to-bed, the MILD. And so, I didn’t understand, but then I have been woken up.

### 6.3.10 Participant 5 – Dream report 2

I had moved into a new house; I was experiencing it like the previous one. And it’s as if I experienced this thing even with this other one, which was more like a mega villa. I was in a very large room that’s almost like an apartment with wide shelves on these windows that almost cover the entire wall, they are very tall. The ceiling is very high. I remember the plants. There are several, both on this shelf, inside and outside. It’s as if the walls were super thick. But there isn’t this perception because of the windows. There were a couple of temporal jumps. But the context was always that of the house there. One is initially, I was in the bedroom. It’s also in the finished house. It’s the moment when I started analyzing more. The windows, the walls, etc. Because there was something strange. I wasn’t aware that it was a dream, but it’s as if something didn’t add up. And for some reason, I ended up outside, as if there was a French door in this room of mine, that leads to the back of the house, on the ground floor. I went out, and there was like a garden, it was mainly a gravel area and a flowerbed that surrounded this small path that encircled the house itself. And it was full of plants. And I knew that they were cultivated by my mom. They were decorative plants: succulents, flowers, etc. After going out, because I was actually looking for something I don’t remember. I went back to the bedroom, and there was a temporal jump, like a flashback when the house had just been finished, with the help of workers. They left us something like a treasure chest, in a playful way. There was a porch on the ground floor, outside the French door in my room. There was this chest, they told us to open it and see what was inside. A kind of gift for the completion of the work. We went to open it. I don’t remember what was inside. The third jump was: The finished house, further in time compared to before, my friends were there. There was this very large trampoline, it was placed on a grid on the ground. We were again at the back of the house, outside. And it is a zone with a recreational area: there’s ping pong. There was another group of guys which didn’t interact with us, initially. We started playing on this trampoline, that caused its supports to fall into the holes and it’s as if the trampoline collapsed, partly got stuck and it made it impossible for us to have fun. We gave up the idea of using it shortly after, even though we moved it out of the grid. And we started playing with the other group that was there. I don’t know if I mentioned it, I was with my friends my long-time friends, from my group. Three of them; then, it’s as if there were others, also the others from this group. the other (group of people), I recognized a friend of a friend but, yes: They weren’t people I was very familiar with. We started playing a game that was based on the card you draw; you modify the rules of the game. But it was physical, it wasn’t simply a board game, you moved, did various activities, etc. The first part of the dream was about exploration. It was like a tree parts, like I saw a temple from point of view but always in the same context. The first part I was feeling weird about the contest, I was not lucid but, it was something not right. For example, I had this big natural, it was my bedroom, and the walls were really tall, but also like, from inside there was a lot of space. In the other two, nothing. There were four groups of characters, one of them I didn’t interact directly. Which were the people who worked to build the house where I’ve stayed in the dream, and the other were my parents. then the other two, it was the group of my old friends, but I can just say that for sure two of them I know there were more. Then the other group were not friends but people that I don’t know well, same age of me and my friends. There were more than one for sure. It was kind of natural, and I was feeling fine. Natural and a lot of fun.

### 6.3.11 Participant 6 – Dream report 1

Did I say anything about Google? I told a story earlier. I meant that there are people in the room where I am now. You asked me before, I heard a voice, I think yours, “what were you dreaming”. I felt like there was a whole group in the room and I was sitting over here. Then I said I need to have a quick drink. Then I went to the water for a minute, I don’t think there’s any water in the room, but I walked to the door, and it had a sink, I drank water there. And then I went back to bed, and I wanted to tell the group and now I am surprised that there is nobody in the room. I told them that I moved into a new flat-share, and it was quite dingy, very alternative, very sympathetic people but a bit dingy. Behind the house they had a big square and there was a big canvas stretched across the square. Google has launched new projectors, has made big advertising for the new devices 3D, they said, the old stuff is no longer needed. Then there was a big presentation on the square in the back with many people which was 3D animated. Before, there was a lot of water, glasses standing there. I went back to bed, and I heard your voice, and I realized that there were no people in the room and there were about five or six people. They were listening intently, as if they were let into this room that night so they could listen to how dreams work. Then I told them about the new WG and the big cloth that is over the place. The projects that Google launched and the show that didn’t happen and people who were on the sides on the cloth and on the cloth, the cloth had the height of a shade cloth of four, three meters and was very strongly stretched. Extra stable on which you can sit on it. There was this demo, this presentation of the new projectors, Hilograms. I dreamt that and first told it to this group of people, before I realized the second time that there was nobody in the room. I have to go over there and have a drink, there were a lot of glasses, also a sink. Then I quickly looked back and saw that a whole group was listening on the sofa. I still had my eyes closed and told them what I had already told them with Google, told them I had to go for a drink and then I opened my eyes and saw that they were all in the room, walked over to the glasses and heard your voice the second time and realized that I hadn’t told them anything yet, hadn’t used my voice at all. I moved into the flat share, I have a girlfriend and I had the feeling that I had to change, move to another place so that it is more fancy and even more running. In this WG I didn’t even have a real place. I had more of a corner where I lived, not a real place, like a pet. The people who lived there were likable, but it was surreal with them. I was in the room once and there was someone in the corner learning which I didn’t even know.

### 6.3.12 Participant 7 – Dream report 1

I was walking home through the city. Then some guy ran past me, all weird and then I thought, yeah easy let’s get out of the way. And afterwards he turns around and then there was another colleague of his and they were pretty drunk, after that they are somehow lying around on the meadow. Afterwards they wanted me to carry their stuff for them.

## 6.4 Appendix D: Online Questionnaire

LD Questionnaire

1. Personal Information: First name:

Lastname.:

Age:

Gender:

Occupation:

- Student
- Retired
- Füll-time employee
- Part-time employee
- Self-employed
- Unemployed
- Others:
- Juggling:

How good would you describe your juggling skills on a scale from 1-5? (1 not good at all, 5 very good)

Can you juggle with three objects? (yes/no)

How confident are you in your juggle skills on a scale from 1-5? (1 not confident at all, 5 extremely confident)

1. Dream Recall:

How often do you remember your dreams in the last few months?

- Almost every morning
- Several times a week
- About once a week
- 2 to 3 times a month
- About once a month
- Less than once a month
- Not at all

1. Lucid Dreaming Experience:

Have you ever experienced a lucid dream before? (yes/no) How frequently da you typically have lucid dreams?

- Rarely (less than once per month)
- Occasionally (1-3 times per month)
- Moderately (1-2 times per week)
- Frequently (3-5 times per week)
- Very frequently (more than 5 times per week)

How do you usually become aware that you are dreaming? (Select all that apply)

- Reality testing
- Dream signs or anomalies
- Intention setting before sleep.
- Spontaneous realization
- others

1. lnduction Techniques:

What specific techniques or methods have you tried to induce lucid dreams? Select all that apply and rate them on a .scale from 1 to 5 (1 not effective at all, 5 extremely effective)

- Reality testing
- Noticing the unreal nature of the dream while dreaming (DILD)
- Intention setting before sleep.
- Wake back to bed (WBTB)
- setting an intention to remember to recognize when one is dreaming (MILD)
- maintaining awareness while transitioning from wakefulness to sleep (WILD)
- Focusing on sensory stimuli and repeatedly shifting the attention between different sensations (SSILD)
- Others

Do you keep a dream journal to record your dreams? {yes/no) How often do you use your dream journal in the last few months?

- Almost every morning
- Several times a week
- About once a week
- 2-3 times a month
- About once a month
- Less than once a month
- Not at all

On a scale from 1-5, how effective has keeping a dream journal been for improving your lucid dreaming skills? (1 not effective at all, S extremely effective)

How frequently do you practice lucid dreaming techniques?

- Rarely (less than once per month)
- Occasionally (1-3 times per month)
- Moderately (1-2 times per week)
- Frequently (3-5 times per week)
- Very frequently (more than 5 times per week)

1. Reality Testing:

Do you perform reality tests during your waking life to check if you are dreaming? (Yes/No)

What methods do you use for reality testing? Select all that apply and rate them on a scale from 1 to 5 (1 not effective at all, S extremely effective)

- searching for abnormal things
- Double-checking the time
- trying to push your finger through your palm.
- check your reflection to see if it looks normal.
- look at your hand
- Reading
- trying to switch the light on/off.
- Dream Goals:

Do you set specific goals or intentions for your dreams7 (yes/no)

On a scale of 1-5, how successful have you been in achieving your dream goals? (1 not successful at all, 5 extremely successful, (1 don’t set goals))

1. Dream Control:

How do you usually behave when you become lucid? (Select all that apply)

- Changing the dream scenery or environment
- Interacting with dream characters
- Manipulating objects or events
- Flying or levitation
- Just observe.
- Other (please specify)

Have you ever attempted to control or manipulate the events in your dreams? (yes/no)

On a scale of 1-5, how would you rate your ability to control your dreams regarding the following items? (1 not .successful at all, 5 extremely successful)

- Own dream body
- Other dream characters’ bodies
- Own dream body movement
- Own actions
- Other dream characters’ actions
- Objects
- Environment
- Gravity

What challenges or limitations have you experienced in controlling aspects of your dreams? (Select all that apply)

- Maintain lucidity.
- Remembering your goals
- loosing focus
- lmagining yourself doing something supernatural
- Having difficulty distinguishing between dreams and real-life situations
- Others

In controlling your dream, what is the challenge with…

…your own dream bodies?

….other dream character’s’ bodies?

….your own dream body movement?

….your own dream actions?

… other dream characters’ actions7

… the objects in your dream?

… the dream environment in your dream?

….time in your dream?

….gravity in your dream?

How much do the challenges you me111tioned above affect the co111trol over your dream? Rate the challenges of each item on a scale from 1-5. (1 - no effect on the dream co111trol at all, 5 - affects the dream control extremely)

What are challenges you already managed to do in a lucid dream? (Select all that apply)

- Communicate with other people in your dream.
- Deliberately shape your environment
- Flying with full control
- Make day turn to night.
- Going through walls, or things
- Going through dream characters
- Eat food.
- Others

1. Motivation and Persistence:

Please rate the following sentences on a scale from 1 ‘I do not agree at all’ to 5’ agree completely’.

- I often come up with new ideas on an older problem or project.
- I remain motivated even with activities that take up several months. 1 have a good ability to focus on daily tasks.
- Long term goals motivate me to tackle day-to-day difficulties.
- Once I decide to do something1 1 don’t give up until I reach the goal. 1 keep thinking of personal goals that I had to give up.
- I purposefully pursue the achievement of projects that I believe in.
- I continue a difficult task even when the others have already given up on it. 1 often find myself thinking about older plans that I have abandoned.
- I invest time and effort in ideas and projects that require years of work and patience.
- The more difficult a task is1 the more determined I am to finish it.
- lt’s hard form to detach from an important project that I have given up in favor of others.

Do you use lucid dreaming for problem solving? (yes/no)

What are your reasons or motivations for lucid dreaming? (Select all that apply)

- For fun
- Social skills
- Creativity
- Wish fulfillment.
- Problem solving
- Emotional regulation
- Doing things, 1 normally can’t do but are possible in real life.
- Overcoming fears/nightmares
- Doing supernatural things
- Training motor skills
- Others

1. Self-Efficacy

How certain are you that you can successfully execute juggling in a lucid dream?

Rate your degree of confidence by recording a number from O to 100 using the scale below. (O = extremely uncertain; 100 = extremely certain)

How certain are you that you can successfully execute juggling in the waking state”?

Rate your degree of confidence by recording a number from O to 100 using the scale below. (O = extremely uncertain; 100 = extremely certain)

Rate your degree of confidence by recording a number from O to 100 using the scale below.

Cannot do at all (0) Moderately can do (50) Highly certain can do (100)

1. Personal Beliefs and Attitudes:

Do you believe you can get lucid in your dream? (yes/no)

Do you think lucid dreaming can have an impact on your waking life? (yes/no) Do you believe you could juggle in a lucid dream? (yes/no)

1. Stress levels:
2. 1. never 2 - almost never 3 - sometimes 4 - fairly often 5 - very often

- In the last month, how often have you been upset because of something that happened unexpectedly?
- In the last month, how often have you felt that you were unable to control the important things in your life?
- In the last month, how often have you felt nervous and stressed?
- In the last month, how often have you felt confident about your ability to handle your personal problems?
- In the last month, how often have you felt that things were going your way?
- In the last month, how often have you found that you could not cope with all the things that you had to do?
- In the last month, how often have you been able to control irritations in your life?
- In the last month, how often have you felt that you were on top of things?
- In the last month, how often have you been angered because of things that happened that were outside of your control?
- In the last month, how often have you felt difficulties were piling up so high that you could not overcome them?
3. 1. Mindfulness:

Scale the following items from 1 - Never or very rarely true, 2 - Rarely true, 3 - Sometimes true, 4 - Often true, 5 - Very often or always true or other way around with the ‘R’ questions (reversed).

- I’m good at finding words to describe my feelings. (D) FFQJM 2
- When I have a sensation in my body, it’s difficult for me to describe it because I can’t find the right words. (D-R) FFQJM 22
- When I’m walking, I deliberately notice the sensations of my body moving. (OBS) FFQM 1
- I notice visual elements in art or nature, such as colors, shapes, textures, or patterns of light and shadow. (OBS) FFQM 31
- I believe some of my thoughts are abnormal or bad and I shouldn’t think that way. (NJ-R) FFQM 14
- I make judgments about whether my thoughts are good or bad. (NJ-R) FFQM 17
- When I have distressing thoughts or images, I “step back’’ and am aware of the thought or image without getting taken over by it. (NR) FFQJM 19
- When I have distressing thoughts or images, I am able just to notice them without reacting. (NR) FFQM 29
- I do jobs or tasks automatically without being aware of what I’m doing. (AA-R) FFQM 34
- I am easily distracted. (AA-R) FFQM 13
- I don’t pay attention to what I’m doing because I’m daydreaming, worrying, or otherwise distracted. (AA-R) fFQM 8

How often do you practice the following mindfulness exercises?

- Meditation

◦ Never
◦ Rarely (less than once per month)
◦ Occasionally (1-3 times per month)
◦ Moderately (1-2 times per week)
◦ Frequently (3-5 times per week)
◦ Very frequently (more than 5 times per week)
- Yoga
- ◦ Never
- ◦ Rarely (less than once per month)
- ◦ Occasionally (1-3 times per month)
- ◦ Moderately (1-2 times per week)
- ◦ Frequently (3-5 times per week)
- ◦ Very frequently (more than 5 times per week)

- Hypnosis

◦ Never
◦ Rarely (less than once per month)
◦ Occasionally (1-3 times per month)
◦ Moderately (1-2 times per week)
◦ Frequently (3-5 times per week)
◦ Very frequently (more than 5 times per week)
- None
- Others

## 6.5 Appendix E: Results of the Online Questionnaire

### 6.5.1 Table E.1 Demographics

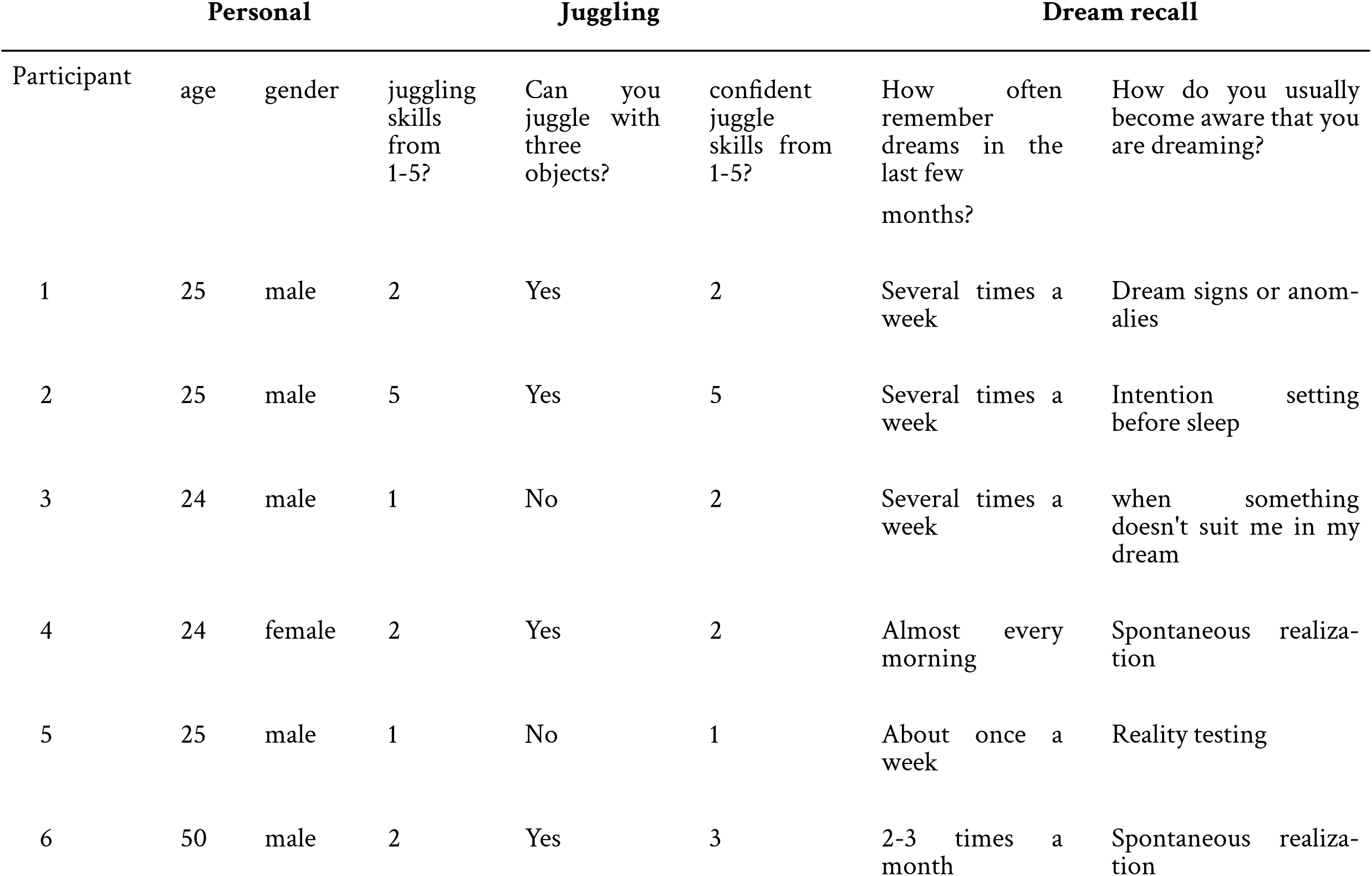
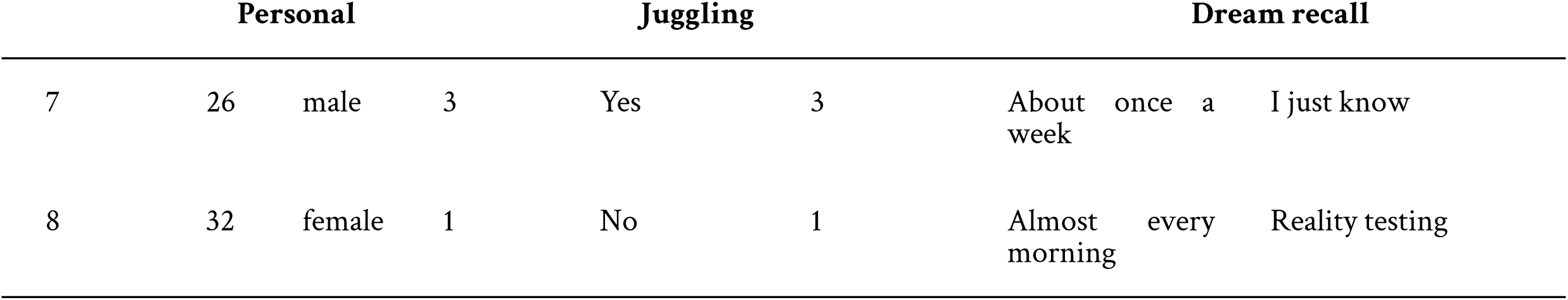

### 6.5.2 Table E.2 Induction techniques

|  | Induction techniques |  |  |  |  |  |  |  |  |  |  |
| --- | --- | --- | --- | --- | --- | --- | --- | --- | --- | --- | --- |
|  | What specific techniques or methods have you tried to induce lucid dreams? Select all that apply and rate them on a scale from 1 to 5 (1 not effective at all, 5 extremely effective) |  |  |  |  |  |  |  |  |  |  |
| Participant | Reality testing | Noticing the unreal nature of the dream while dreaming (DILD) | Intention setting before sleep | Wake back to bed (WBTB) | Setting an intention to remember to recognize when one is dreaming (MILD) | Maintaining awareness while transitioning from wakefulness to sleep (WILD) | Focusing on sensory stimuli and repeatedly shifting the attention between different sensations (SSILD) | Do you keep a dream journal to record your dreams? | How often do you use your dream journal in the last few months? | On a scale from 1-5, how effective has keeping a dream journal been for improving your lucid dreaming skills? (1 not effective at all, 5 extremely effective) | How frequently do you practice lucid dreaming techniques? |
| 1 | No | No | No | Yes (Similar to my clocks go off) | No | No | No | No |  |  | Frequently (3-5 times per week) |
| 2 | Yes | Yes | Yes | Yes | No | Yes | Yes | Yes | About once a month | 4 | Occasionally (1-3 times per month) |
| 3 | Yes | Yes (4) | No | No | No | No | No | Yes | 2-3 times a month | 3 | Occasionally (1-3 times per month) |
| 4 | Yes (2) | No | Yes (2) | Yes (3) | No | Yes (3) | No | Yes | Less than once a month | 2 | Rarely (less than once per month) |
| 5 | Yes (5) | No | Yes (2) | Yes (4) | Yes (4) | Yes (3) | No | No |  |  | Rarely (less than once per month) |
| 6 | No | Yes | No | No | No | No | No | No |  |  | Rarely (less than once per month) |
| 7 | No | Yes (5) | No | No | No | No | No | Yes | Less than once a month | 3 | Rarely (less than once per month) |
| 8 | Yes (5) | No | No | Yes (4) | No | No | No | Yes | 2-3 times a month | 4 | Occasionally (1-3 times per month) |

### 6.5.3 Table E.3 Reality testing

| Participant | Do you perform reality tests during your waking life to check if you are dreaming? | Searching for abnormal things | Double-checking the time | Trying to push your finger through your palm | Check your reflection to see if it looks normal | Look at your hands | Reading | Trying to switch the light on/off | Other |
| --- | --- | --- | --- | --- | --- | --- | --- | --- | --- |
| 1 | 0 | 0 | 0 | 0 | 0 | 0 | 0 | 0 | Testing the reality with my talisman |
| 2 | Yes | Yes | 0 | 0 | 0 | Yes | 0 | 0 |  |
| 3 | 0 | Yes (4) | 0 | 0 | 0 | 0 | 0 | 0 |  |
| 4 | Yes | Yes (2) | 0 | 0 | 0 | Yes (2) | 0 | 0 |  |
| 5 | 0 | 0 | 0 | 0 | 0 | Yes | 0 | 0 |  |
| 6 | 0 | Yes (2) | 0 | 0 | 0 | 0 | 0 | 0 |  |

| What methods do you use for reality testing? Select all that apply and rate them on a scale from 1 to 5 (1 not effective at all, 5 extremely effective) |  |  |  |  |  |  |  |  |  |
| --- | --- | --- | --- | --- | --- | --- | --- | --- | --- |
| 7 | 0 | 0 | 0 | 0 | 0 | 0 | 0 | 0 | Trying to fly (5) |
| 8 | Yes | Yes (4) | 0 | 0 | 0 | 0 | 0 | Yes (5) |  |

### 6.5.4 Table E.4 Dream goals

| Dream goals |  |  |
| --- | --- | --- |
| Participant | Do you set specific goals or intentions for your dream? | how successful have you been? (1 not successful at all, 5 extremely successful) |
| 1 | Yes | 4 |
| 2 | No |  |
| 3 | No |  |
| 4 | Yes | 2 |
| 5 | No |  |
| 6 | No |  |
| 7 | No |  |
| 8 | Yes | 3 |

### 6.5.5 Table E.5.1 Dream control 1

How do you usually behave when you become lucid? (Select all that apply)
| Participant | Changing the dream scenery or environment | Interacting with dream characters | Manipulating objects or events | Flying or levitation | Just observe | Other |
| --- | --- | --- | --- | --- | --- | --- |
| 1 | Yes | Yes | Yes | Yes | Yes | Create a world starting with just a white blanket room |
| 2 | No | No | No | Yes | Yes |  |
| 3 | Yes | Yes | No | No | Yes |  |
| 4 | Yes | No | No | Yes | Yes |  |
| 5 | No | No | No | Yes | No |  |
| 6 | Yes | Yes | No | No | No |  |
| 7 | No | No | No | No | Yes |  |
| 8 | Yes | No | Yes | Yes | No |  |

### 6.5.6 Table E.5.2 Dream control 2

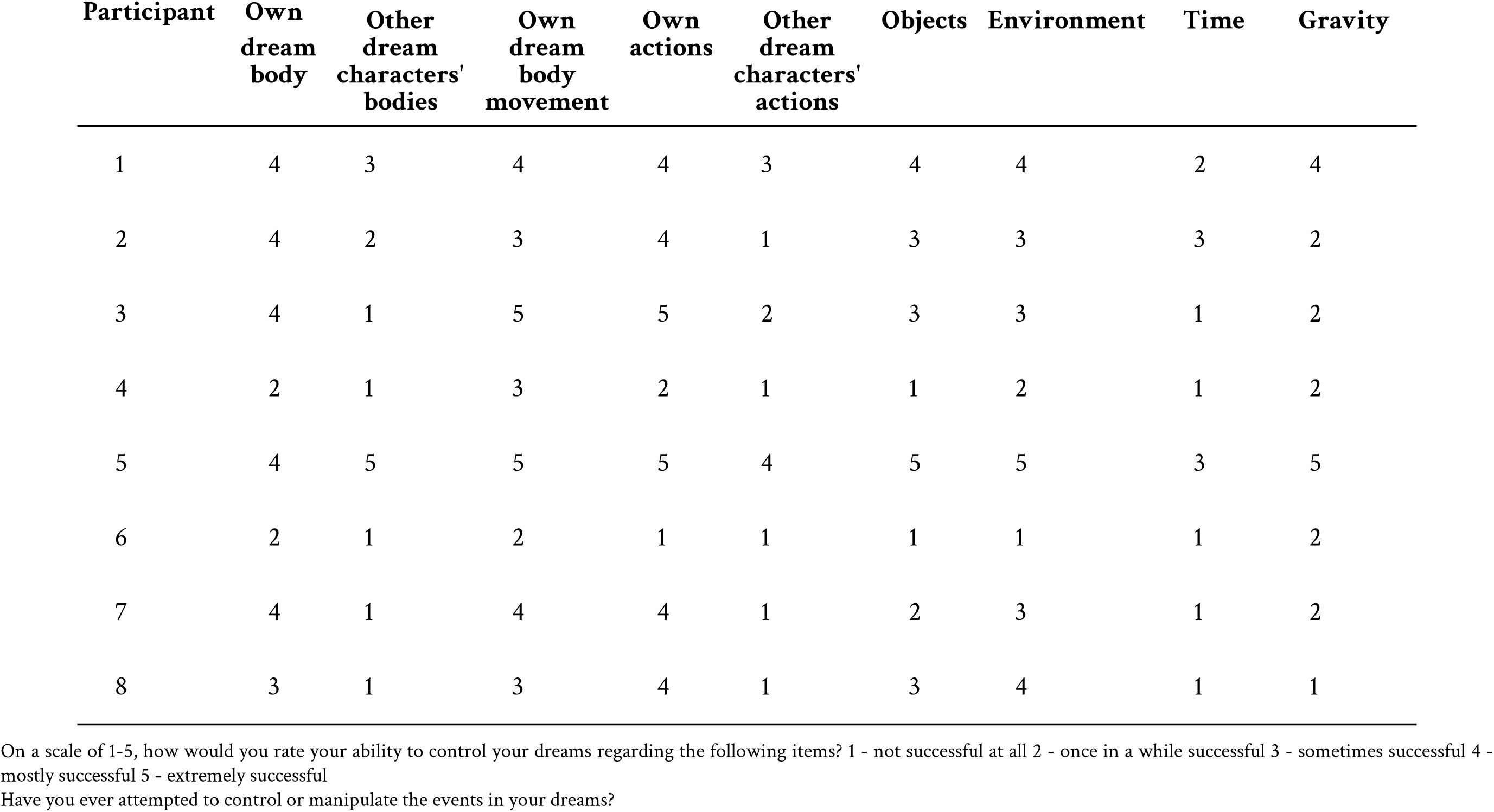

### 6.5.7 Table E.5.3 Dream control 3

| Participant | Maintaining<br>Lucidity | Remembering<br>your goals | Letting<br>supernatural<br>things hap-<br>pen | Loosing<br>focus | Imagining<br>yourself<br>doing some-<br>thing super-<br>natural | Having difficulty<br>distinguishing<br>between dreams<br>and real-life situa-<br>tions | Other |
| --- | --- | --- | --- | --- | --- | --- | --- |
| 1 | Yes | Yes | Yes | Yes | Yes | Yes |  |
| 2 | Yes | No | Yes | No | Yes | No |  |
| 3 | Yes | No | Yes | No | No | Yes |  |
| 4 | Yes | No | No | Yes | No | No |  |
| 5 | No | Yes | No | No | No | No | I eas-<br>ily<br>wake<br>up |
| 6 | No | No | No | Yes | No | No |  |
| 7 | No | Yes | No | No | No | No |  |
| 8 | No | No | Yes | No | No | No |  |

What challenges or limitations have you experienced in controlling aspects of your dreams? (Select all that apply)

### 6.5.8 Table E.5.4 Dream control 4

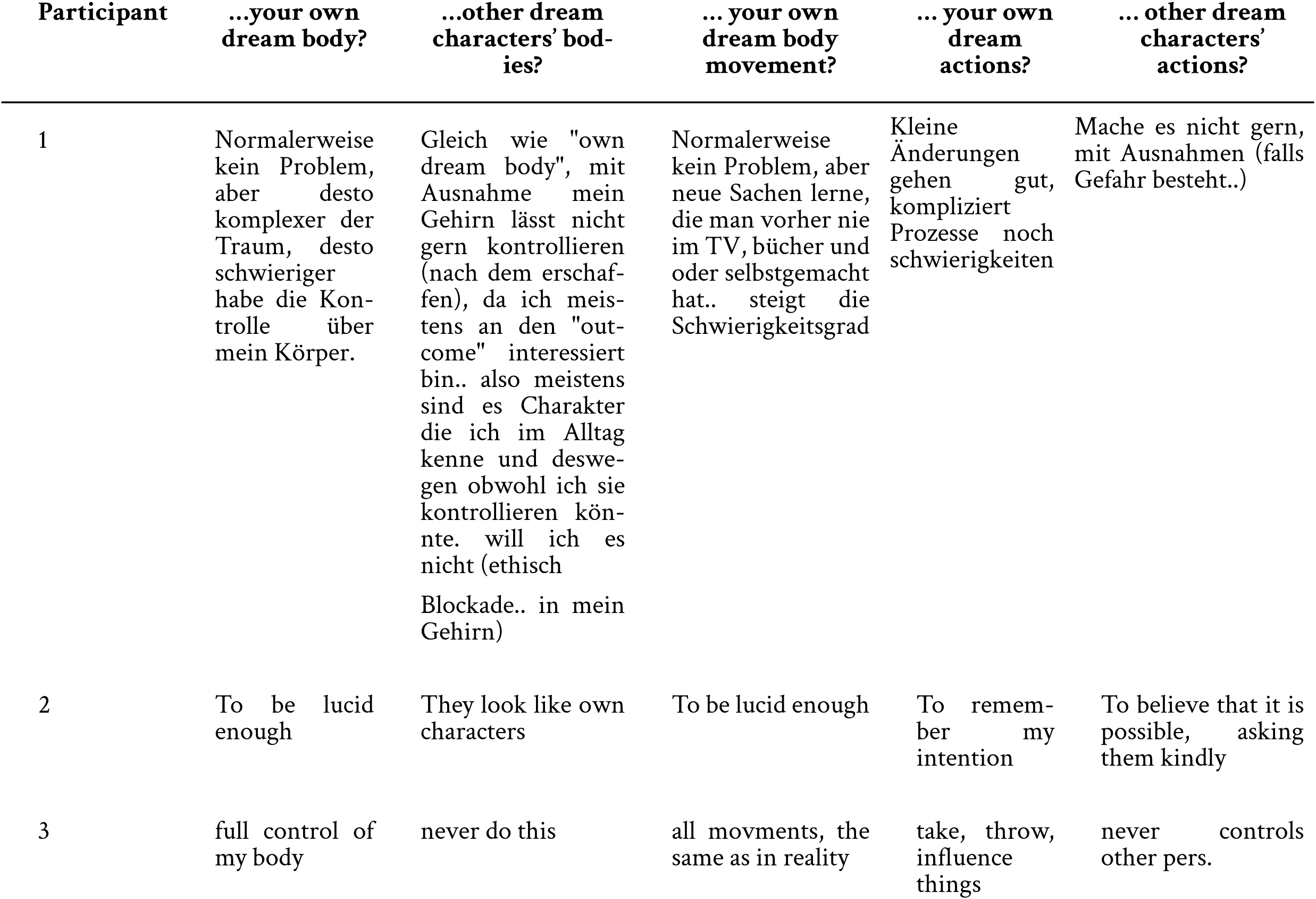
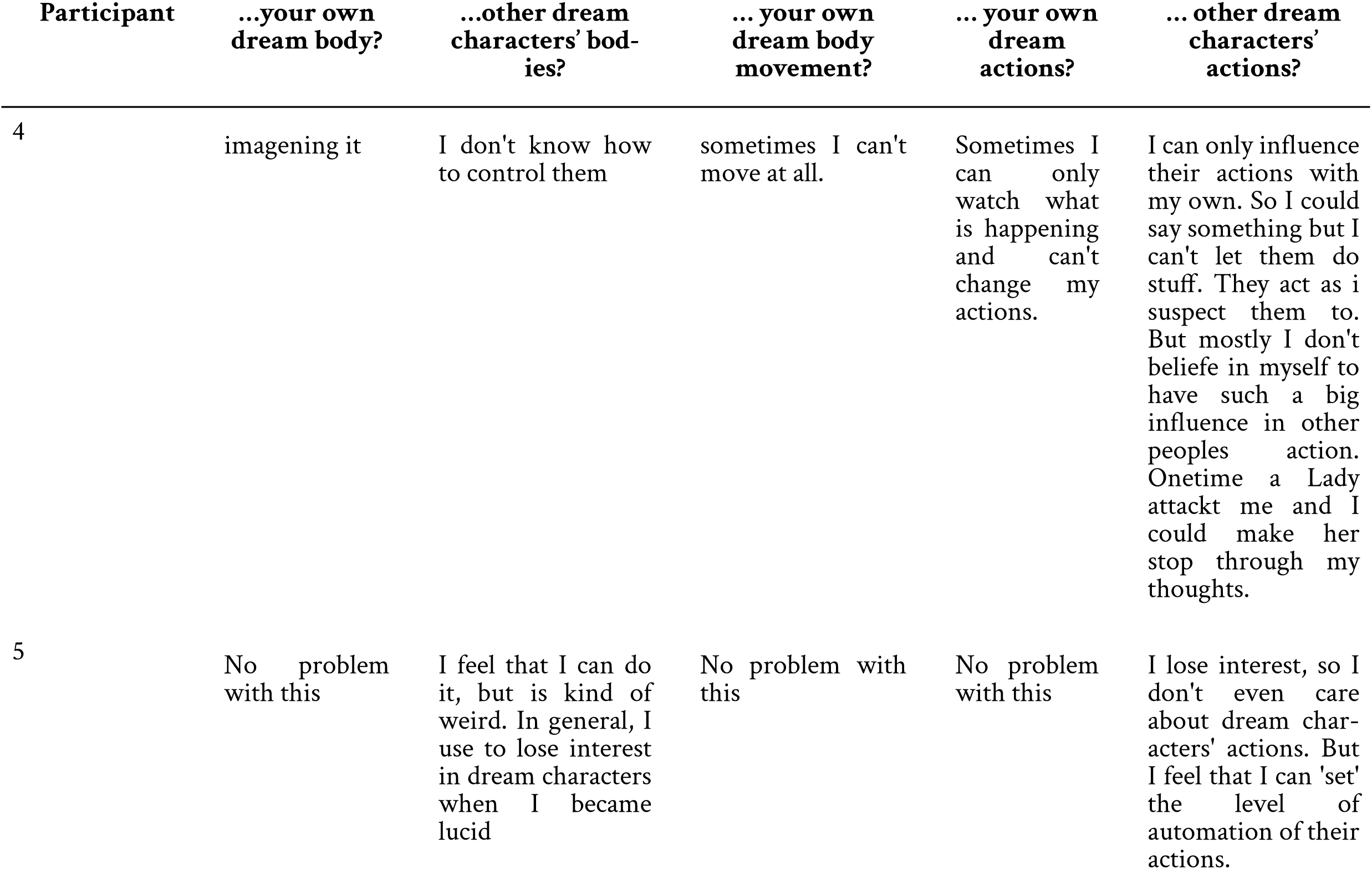
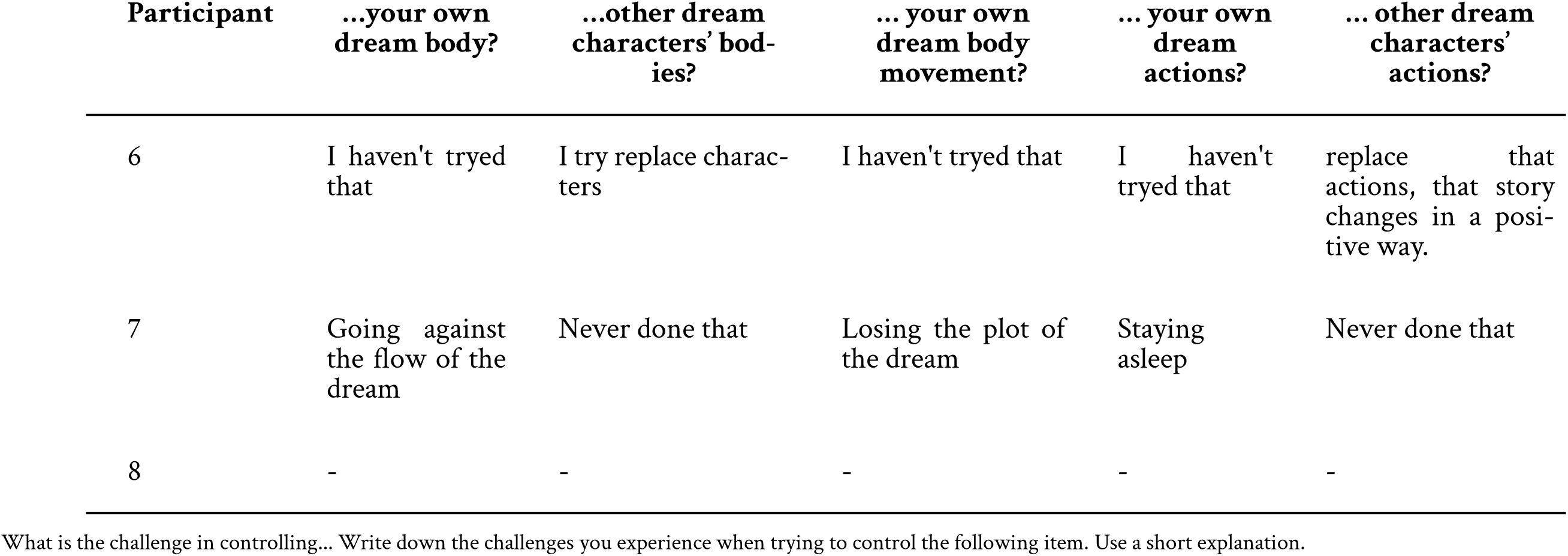

### 6.5.9 Table E.5.5 Dream control 5

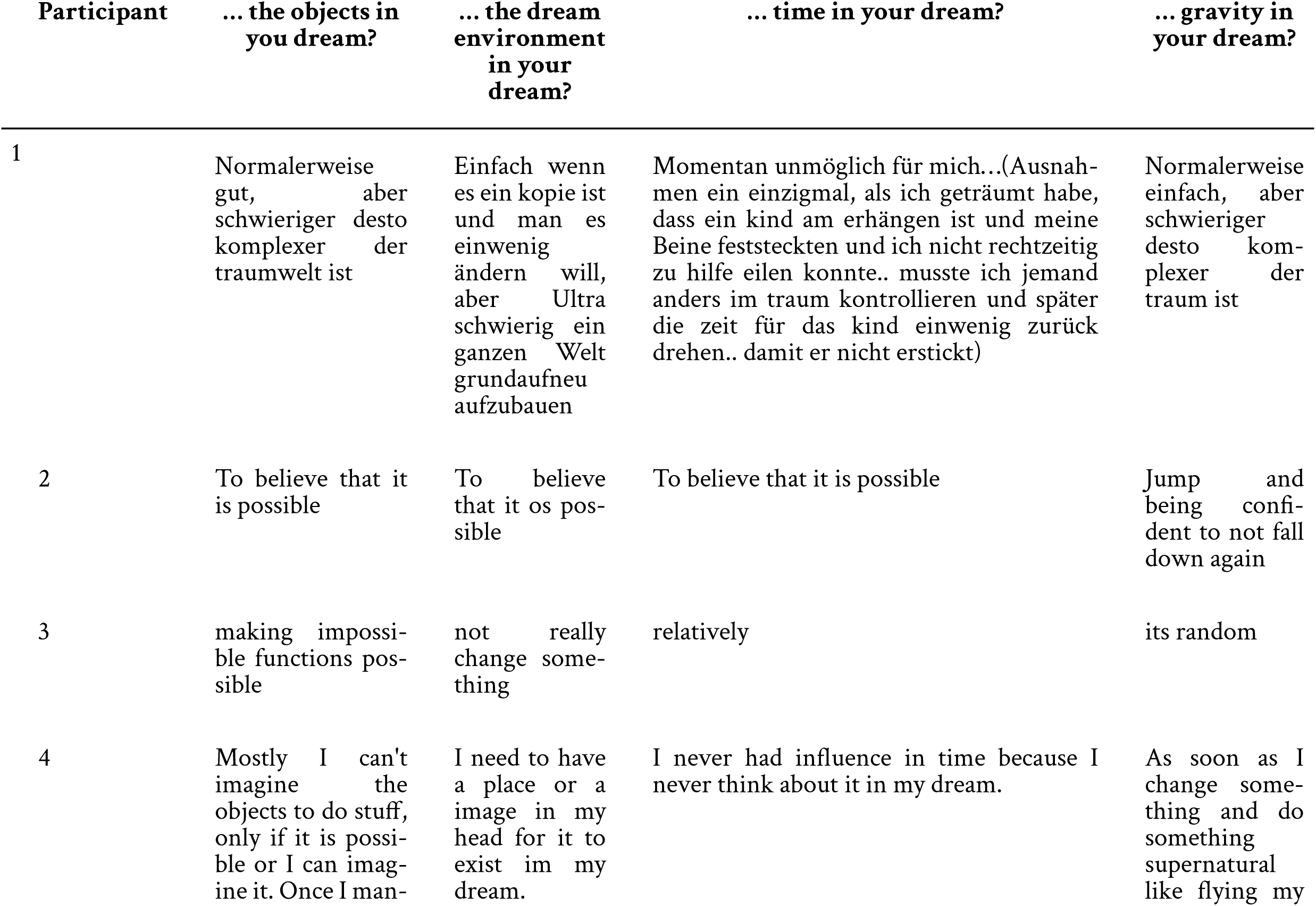
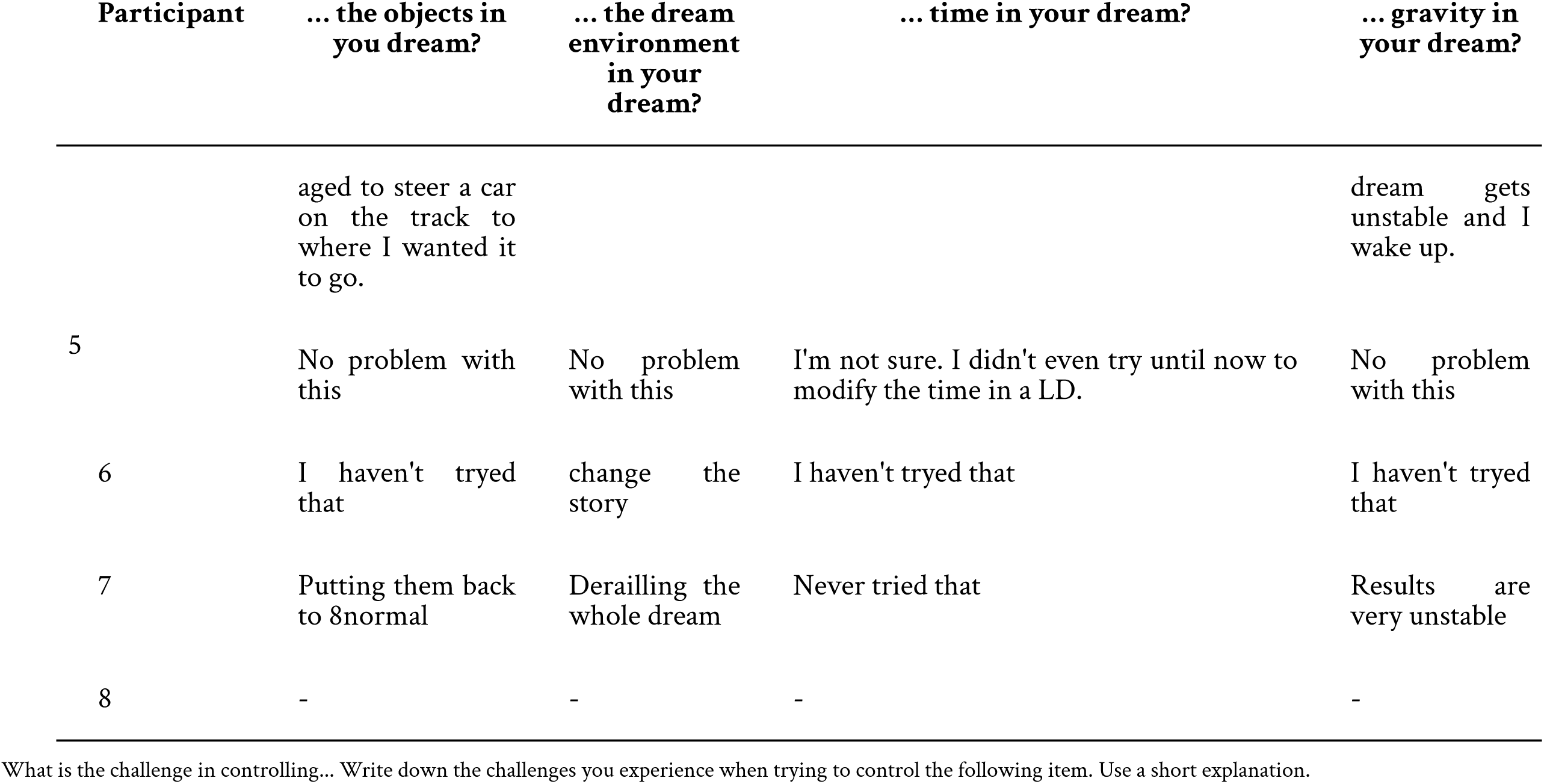

### 6.5.10 Table E.5.6 Dream control 6

| Participant | your own dream body | other dream characters' bodies | your own dream body movement | own dream actions | other dream characters' actions | the objects in your dream | the dream environment in your dream | time in your dream | gravity in your dream |
| --- | --- | --- | --- | --- | --- | --- | --- | --- | --- |
| 1 | 1 | 1 | 1 | 1 | 1 | 2 | 2 | 5 | 4 |
| 2 | 5 | 3 | 5 | 5 | 3 | 5 | 4 | 3 | 5 |
| 3 | 1 | 1 | 2 | 5 | 3 | 4 | 5 | 1 | 2 |
| 4 | 2 | 1 | 4 | 4 | 1 | 2 | 2 | 1 | 4 |
| 5 | 1 | 2 | 1 | 1 | 2 | 1 | 1 | 4 | 1 |
| 6 | 1 | 1 | 1 | 3 | 1 | 2 | 3 | 1 | 1 |
| 7 | 5 | 1 | 4 | 4 | 1 | 4 | 1 | 1 | 3 |
| 8 | 1 | 1 | 1 | 1 | 1 | 1 | 1 | 1 | 1 |

How much do the challenges you mentioned above affect the control over your dream? Rate the challenges for each item on a scale from 1-5. (1 - no effect on the dream control at all, 5 - affects the dream control extremely) the challenge with...

### 6.5.11 Table E.5.7 Dream control 7

| Participant | Communicate with other people in your dream | Deliberately shape your environment | Flying with full control | Make day turn to night | Going through walls or things | Going through dream characters | Other |
| --- | --- | --- | --- | --- | --- | --- | --- |
| 1 | Yes | Yes | Yes | Yes | No | No |  |
| 2 | Yes | No | Yes | Yes | Yes | No | Transform into animals |
| 3 | Yes | No | Yes | Yes | No | Yes |  |
| 4 | Yes | Yes | Yes | No | No | No |  |
| 5 | No | Yes | Yes | No | No | No |  |
| 6 | Yes | No | No | No | No | No |  |
| 7 | Yes | Yes | Yes | No | No | No |  |
| 8 | No | Yes | Yes | No | Yes | No |  |

What are challenges you already managed to do in a lucid dream? (Select all that apply)

### 6.5.12 Table E.6.1 Motivation and persistence 1

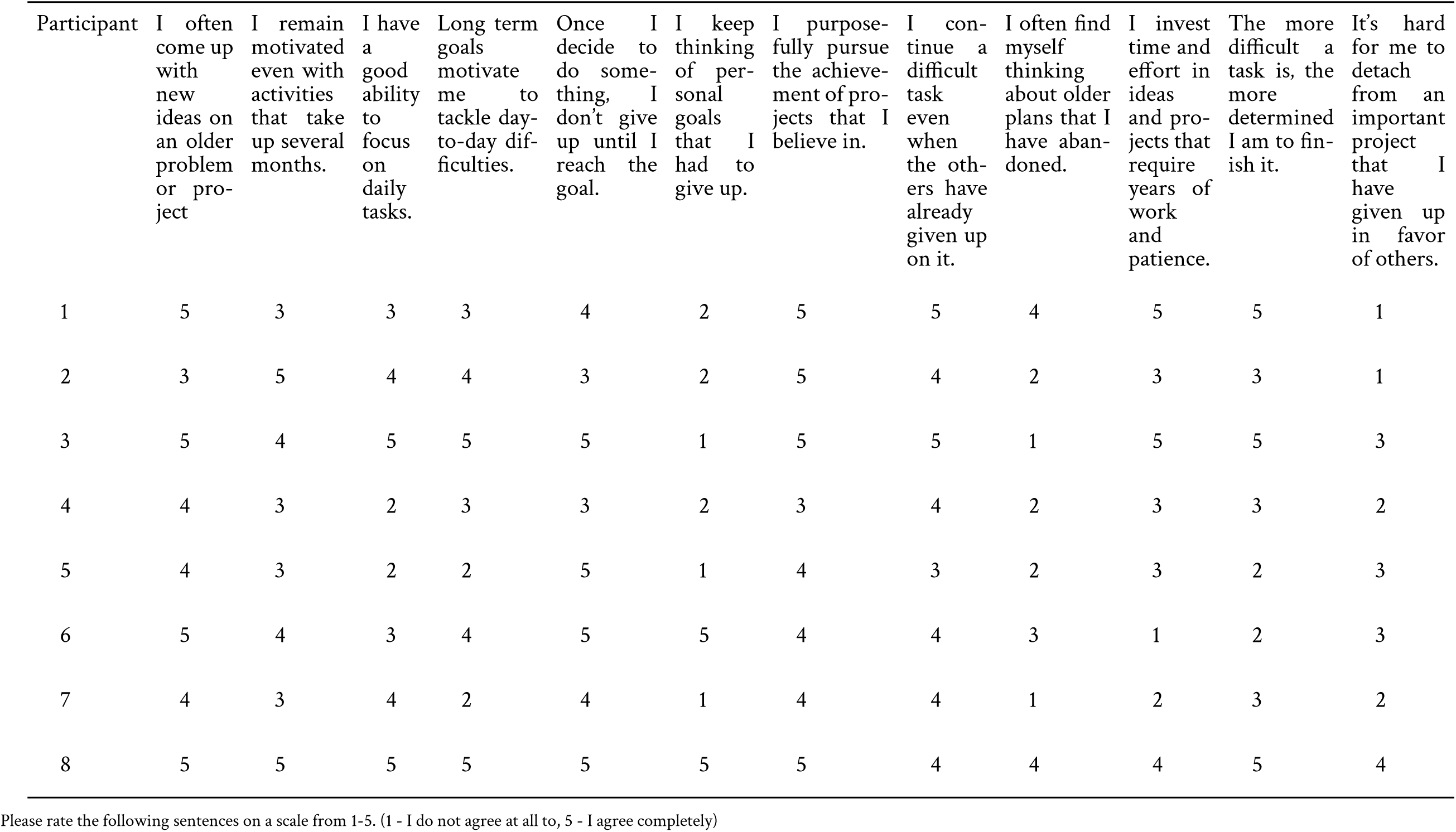

### 6.5.13 Table E.6.2 Motivation and persistence 2

| Participant | Do you use lucid dreaming for problem solving? | For fun | Social skills | Creativity | Wish fulfillment | Problem solving | Emotional regulation | Doing things I normally can't do but are possible in real life | Overcoming fears/nightmares | Doing super-natural things | Training motor skills | Other |
| --- | --- | --- | --- | --- | --- | --- | --- | --- | --- | --- | --- | --- |
| 1 | Yes | Yes | No | No | No | Yes | No | Yes | No | No | No |  |
| 2 | No | Yes | No | Yes | No | No | No | No | No | Yes | No | Doing things, I can't do in real life |
| 3 | No | Yes | Yes | Yes | No | No | No | No | No | Yes | Yes |  |
| 4 | No | Yes | No | Yes | No | No | No | Yes | No | Yes | No |  |
| 5 | No | Yes | No | Yes | No | No | Yes | No | No | Yes | No |  |
| 6 | No | No | No | No | No | No | Yes | No | No | No | No |  |
| 7 | No | Yes | No | No | No | No | No | No | No | No | No |  |
| 8 | Yes | Yes | No | Yes | No | Yes | No | Yes | No | Yes | No |  |

What are your reasons or motivations for lucid dreaming? (Select all that apply)

### 6.5.14 Table E.7 Self-efficacy & personal beliefs and attitudes

| Participant | how certain are you that you can successfully execute juggling in a lucid dream? | how certain are you that you can successfully execute juggling in the waking state? | Do you believe you can get lucid in your dream? | Do you think dreaming can have an impact on your life? | Do you believe you could juggle in a lucid dream? |
| --- | --- | --- | --- | --- | --- |
| 1 | 100 | 50 | Yes | Yes | Yes |
| 2 | 100 | 100 | Yes | Yes | Yes |
| 3 | 40 | 20 | Yes | Yes | Yes |
| 4 | 60 | 20 | Yes | Yes | Yes |
| 5 | 70 | 10 | Yes | Yes | Yes |
| 6 | 30 | 80 | Yes | No | Yes |
| 7 | 50 | 75 | Yes | No | Yes |
| 8 | 50 | 50 | Yes | Yes | Yes |

### 6.5.15 Table E.8 Perceived stress scale

| Participant | ...been upset because of something that happened unexpectedly? | ...felt that you were unable to control the important things in your life? | ...felt nervous and stressed? | ...felt confident about your ability to handle your personal problems? | ...felt that things were going your way? | ...found that you could not cope with all the things that you had to do? | ...been able to control irritations in your life? | ...felt that you were on top of things? | ...been angered because of things that happened that were outside of your control? | ...felt difficulties were piling up so high that you could not overcome them? |
| --- | --- | --- | --- | --- | --- | --- | --- | --- | --- | --- |
| 1 | 0 | 3 | 3 | 1 | 3 | 3 | 2 | 3 | 1 | 0 |
| 2 | 0 | 1 | 2 | 0 | 2 | 2 | 2 | 1 | 1 | 1 |
| 3 | 0 | 0 | 0 | 0 | 1 | 1 | 1 | 0 | 0 | 0 |
| 4 | 1 | 1 | 2 | 1 | 1 | 2 | 1 | 2 | 1 | 2 |
| 5 | 1 | 3 | 3 | 3 | 2 | 4 | 1 | 3 | 1 | 1 |
| 6 | 3 | 2 | 3 | 2 | 2 | 2 | 1 | 2 | 2 | 2 |
| 7 | 2 | 1 | 2 | 0 | 1 | 1 | 3 | 1 | 3 | 1 |
| 8 | 3 | 1 | 2 | 1 | 1 | 3 | 1 | 1 | 2 | 1 |

In the last month, how often have you...

### 6.5.16 Table E.9.1 Mindfulness 1

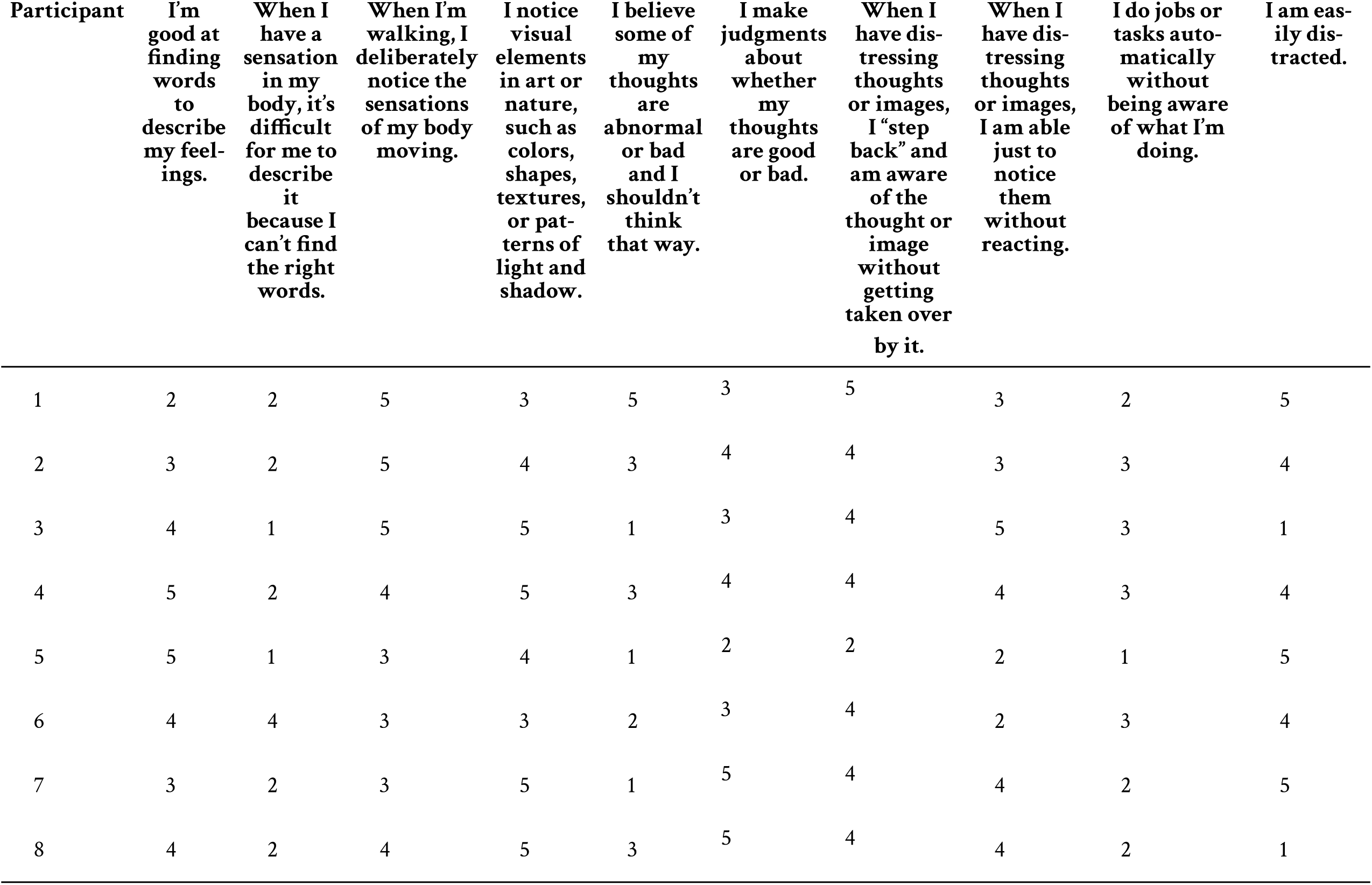

| Participant | I'm good at finding words to describe my feelings. | When I have a sensation in my body, it's difficult for me to describe it because I can't find the right words. | When I'm walking, I deliberately notice the sensations of my body moving. | I notice visual elements in art or nature, such as colors, shapes, textures, or patterns of light and shadow. | I believe some of my thoughts are abnormal or bad and I shouldn't think that way. | I make judgments about whether my thoughts are good or bad. | When I have distressing thoughts or images, I "step back" and am aware of the thought or image without getting taken over by it. | When I have distressing thoughts or images, I am able just to notice them without reacting. | I do jobs or tasks automatically without being aware of what I'm doing. | I am easily distracted. |
| --- | --- | --- | --- | --- | --- | --- | --- | --- | --- | --- |
| 1 | 2 | 2 | 5 | 3 | 5 | 3 | 5 | 3 | 2 | 5 |
| 2 | 3 | 2 | 5 | 4 | 3 | 4 | 4 | 3 | 3 | 4 |
| 3 | 4 | 1 | 5 | 5 | 1 | 3 | 4 | 5 | 3 | 1 |
| 4 | 5 | 2 | 4 | 5 | 3 | 4 | 4 | 4 | 3 | 4 |
| 5 | 5 | 1 | 3 | 4 | 1 | 2 | 2 | 2 | 1 | 5 |
| 6 | 4 | 4 | 3 | 3 | 2 | 3 | 4 | 2 | 3 | 4 |
| 7 | 3 | 2 | 3 | 5 | 1 | 5 | 4 | 4 | 2 | 5 |
| 8 | 4 | 2 | 4 | 5 | 3 | 5 | 4 | 4 | 2 | 1 |

### 6.5.17 Table E.9.2 Mindfulness 2

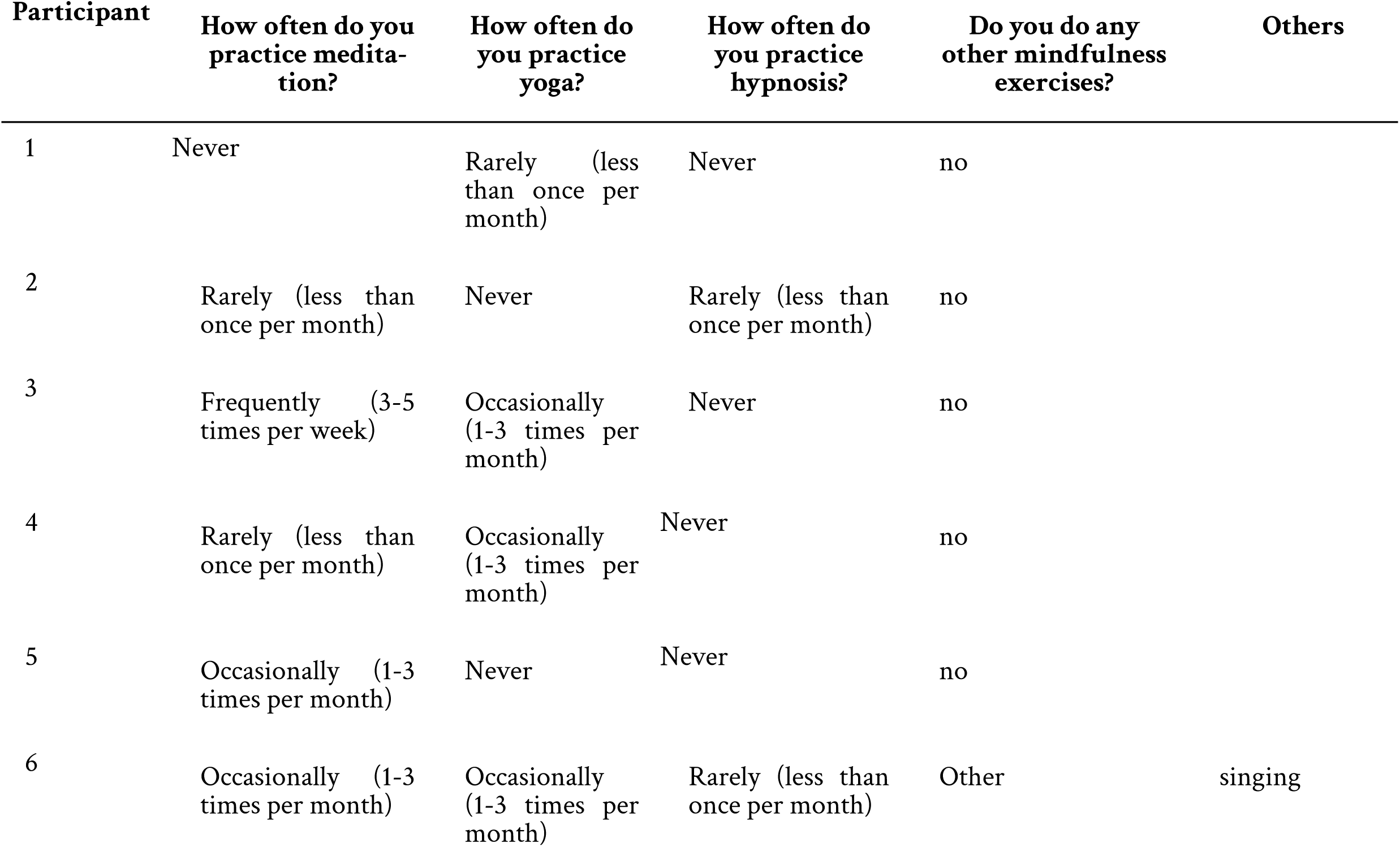
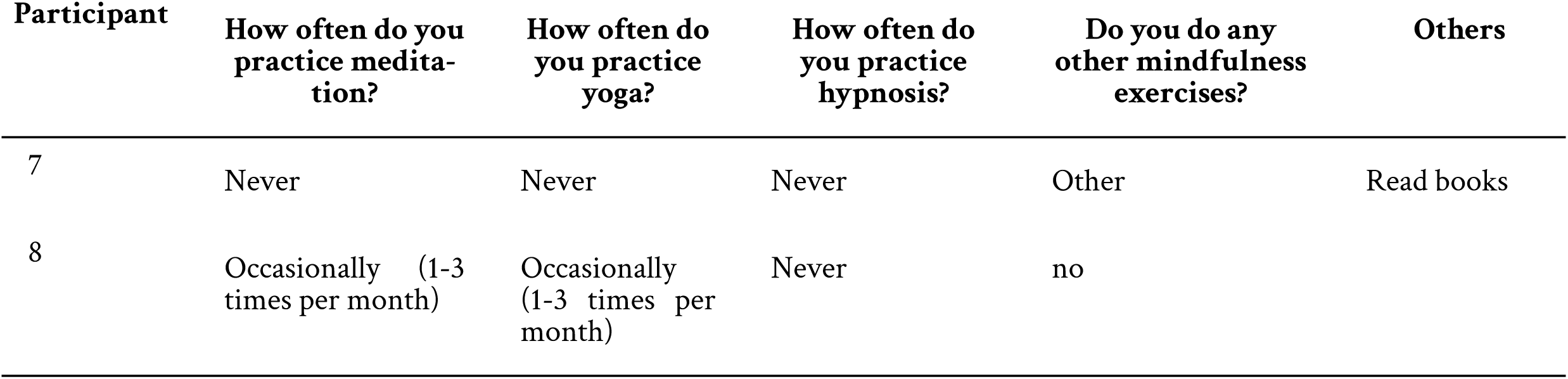

| Participant | How often do you practice meditation? | How often do you practice yoga? | How often do you practice hypnosis? | Do you do any other mindfulness exercises? | Others |
| --- | --- | --- | --- | --- | --- |
| 1 | Never | Rarely (less than once per month) | Never | no |  |
| 2 | Rarely (less than once per month) | Never | Rarely (less than once per month) | no |  |
| 3 | Frequently (3-5 times per week) | Occasionally (1-3 times per month) | Never | no |  |
| 4 | Rarely (less than once per month) | Occasionally (1-3 times per month) | Never | no |  |
| 5 | Occasionally (1-3 times per month) | Never | Never | no |  |
| 6 | Occasionally (1-3 times per month) | Occasionally (1-3 times per month) | Rarely (less than once per month) | Other | singing |

| **Participant** | **How often do you practice meditation?** | **How often do you practice yoga?** | **How often do you practice hypnosis?** | **Do you do any other mindfulness exercises?** | **Others** |
| --- | --- | --- | --- | --- | --- |
| 7 | Never | Never | Never | Other | Read books |
| 8 | Occasionally (1-3 times per month) | Occasionally (1-3 times per month) | Never | no |  |

